# Chemoproteomic study uncovers HemK2/KMT9 as a new target for NTMT1 bisubstrate inhibitors

**DOI:** 10.1101/2021.04.13.439666

**Authors:** Dongxing Chen, Ying Meng, Dan Yu, Nicholas Noinaj, Xiaodong Cheng, Rong Huang

## Abstract

Understanding the selectivity of methyltransferase inhibitors is important to dissect the functions of each methyltransferase target. From this perspective, here we report a chemoproteomic study to profile the selectivity of a potent protein N-terminal methyltransferase 1 (NTMT1) bisubstrate inhibitor NAH-C3-GPKK (*K*_i, app_ = 7±1 nM) in endogenous proteomes. First, we describe the rational design, synthesis, and biochemical characterization of a new chemical probe **6**, a biotinylated analogue of NAH-C3-GPKK. Next, we systematically analyze protein networks that may selectively interact with the biotinylated probe **6** in concert with the competitor NAH-C3-GPKK. Besides NTMT1, the designated NTMT1 bisubstrate inhibitor NAH-C3-GPKK was found to also potently inhibit a methyltransferase complex HemK2-Trm112 (also known as KMT9-Trm112), highlighting the importance of systematic selectivity profiling. Furthermore, this is the first potent inhibitor for HemK2/KMT9 reported to date. Thus, our studies lay the foundation for future efforts towards the development of selective inhibitors for NTMT1 and HemK2/KMT9.

**TOC:** 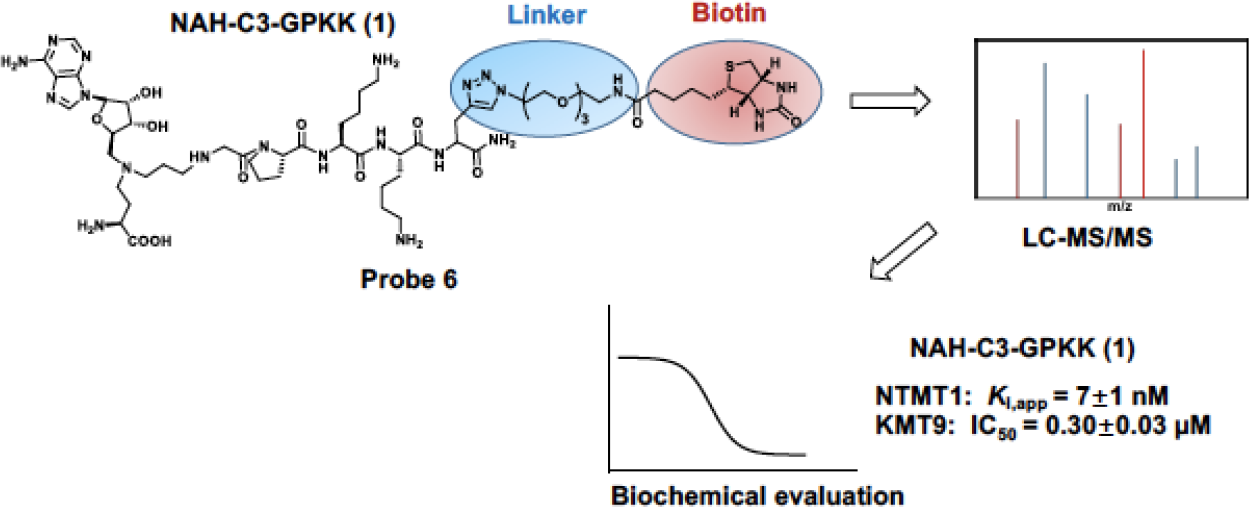

## INTRODUCTION

Protein methylation regulates numerous biological processes, including transcription, DNA repair, and RNA metabolism.^1–3^ Protein methyltransferases (MTs) are the enzymes that catalyze the transfer of a methyl group from the cofactor S-adenosyl-L-methionine (SAM) onto protein substrates. The substrate preference of MTs determines their respective functions and annotations.^4–10^ For example, protein N-terminal MTs (NTMTs), lysine MTs (PKMTs), and arginine MTs (PRMTs) catalyze the methylation on protein α-N-termini, ε-amino group of the lysine, and guanidinium nitrogen atoms of the arginine, respectively.^4–10^ As the dysregulation of MTs has been implicated in various diseases, selectivity of the inhibitors is essential to elucidate the physiological and pharmacological functions of the MTs.^1–3, 11–13^

NTMT1 methylates the protein α-N-terminal amines of a specific motif X-P-K/R, where X represents any amino acid other than D/E.^5, 8, 14^ Substrates of NTMT1 include centromere H3 variants CENP-A/B, the regulator of chromatin condensation 1 (RCC1), damaged DNA-binding protein 2, poly(ADP-ribose) polymerase 3, and transcription factor-like protein MRG15.^5, 15–18^ Recent progress has revealed the important role of NTMT1 in chromosome segregation, mitosis, and DNA damage repair.^18–20^ Knockdown of NTMT1 sensitized the breast cancer cell lines to etoposide and γ irradiation treatment, and knockout of NTMT1 resulted in developmental defects and premature aging in mice.^20, 21^ Therefore, such critical cellular processes highlight the need for the development of highly selective chemical probes for NTMT1. Towards this goal, we have previously developed a series of potent NTMT1 bisubstrate inhibitors by covalently linking the SAM mimic with a peptide substrate to allow them to simultaneously occupy neighboring binding sites of SAM and substrate. Two representative examples are NAH-C3-PPKRIA (*K*_i, app_ = 39±10 nM) and NAH-C4-GPKRIA (*K*_i, app_ = 0.13±0.04 nM) (Figure 1), where NAH is a SAM mimic by replacing the sulfonium ion with a nitrogen atom. Meanwhile, they displayed selectivity over two representative PKMTs (G9a and SETD7) and three PRMTs (PRMT1, 3, and 7) in the biochemical assays.^22–25^ Furthermore, NAH-C4-GPKRIA exhibits selective for NTMT1 over its closely related homolog NTMT2, suggesting the unique active site of NTMT1 and high selectivity of NTMT1 bisubstrate inhibitors.^25^ While NTMT1 bisubstrate inhibitors exhibit decent selectivity among multiple MTs against isolated enzymes in biochemical assays, there is no information on any other targets that those inhibitors may interact within cellular contexts to understand any off-target engagement and delineate the mechanism of action.

**Figure 1.**
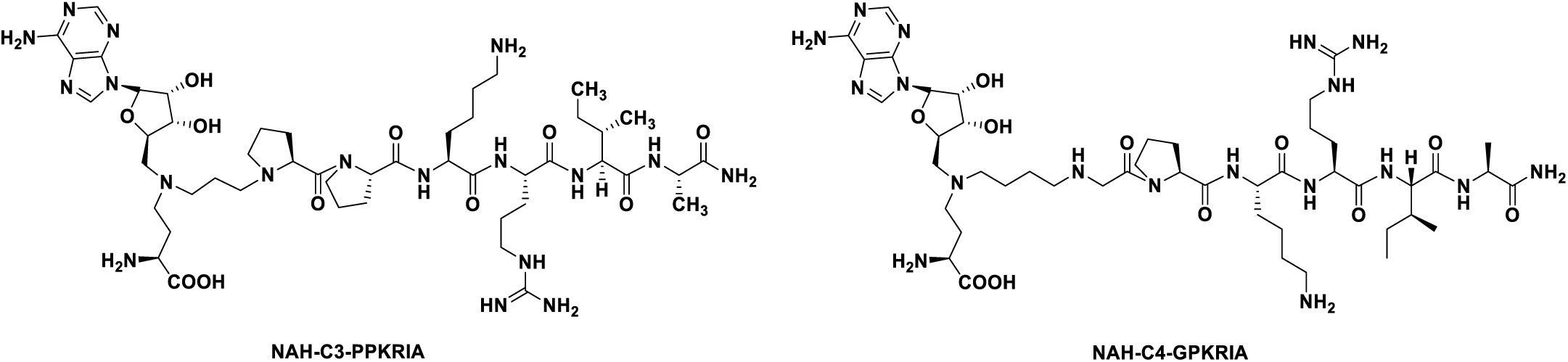
Chemical structures of NTMT1 bisubstrate inhibitors.

Chemoproteomic profiling is a powerful approach to provide a global insight into the cellular targets of an inhibitor in a complex biological environment by utilizing immobilized or photoreactive inhibitors. Here, we design and synthesize a biotinylated derivative of NTMT1 bisubstrate inhibitor **1** (NAH-C3-GPKK) to generate probe **6** (NAH-C3-GPKK-biotin), allowing it to be immobilized with streptavidin beads. Pulldown study of probe **6** confirms its ability to efficiently enrich NTMT1 from the whole cell lysates. In addition, probe **6** also binds to another protein methyltransferase, i.e. HemK2-Trm112 complex (also known as KMT9-Trm112), that methylates both lysines of histone H4 and glutamine of the eukaryotic release factor 1 (eRF1).^4, 26^ Subsequent validation confirms that compound **1** acts as a dual potent inhibitor for NTMT1 and HemK2/KMT9. To our knowledge, this is the first inhibitor reported for HemK2/KMT9, which offers a new revenue for the use of this compound.

## RESULTS AND DISCUSSION

### Design and synthesis

To design the optimal probe for the selectivity profiling of NTMT1 bisubstrate inhibitor, we first investigate the structure-activity relationship (SAR) of the NTMT1 bisubstrate inhibitors. Thus, we synthesized new bisubstrate analogues **1**-**4** with different peptide sequences and variable linkers to choose the optimal position to install a biotin group. Because of its high potency and selectivity, NAH-C4-GPKRIA was selected as our lead compound. According to the co-crystal structures of NTMT1 in complex with previous NTMT1 bisubstrate inhibitors,^24, 25^ we kept the tripeptide GPK intact in our design. The proposed **1**-**4** were synthesized in a convergent manner by reacting the primary amines (**7-8**) with α-bromo peptide on resin and followed by acid treatment as described before (Schemes 1).^23, 25^

**Scheme 1.**
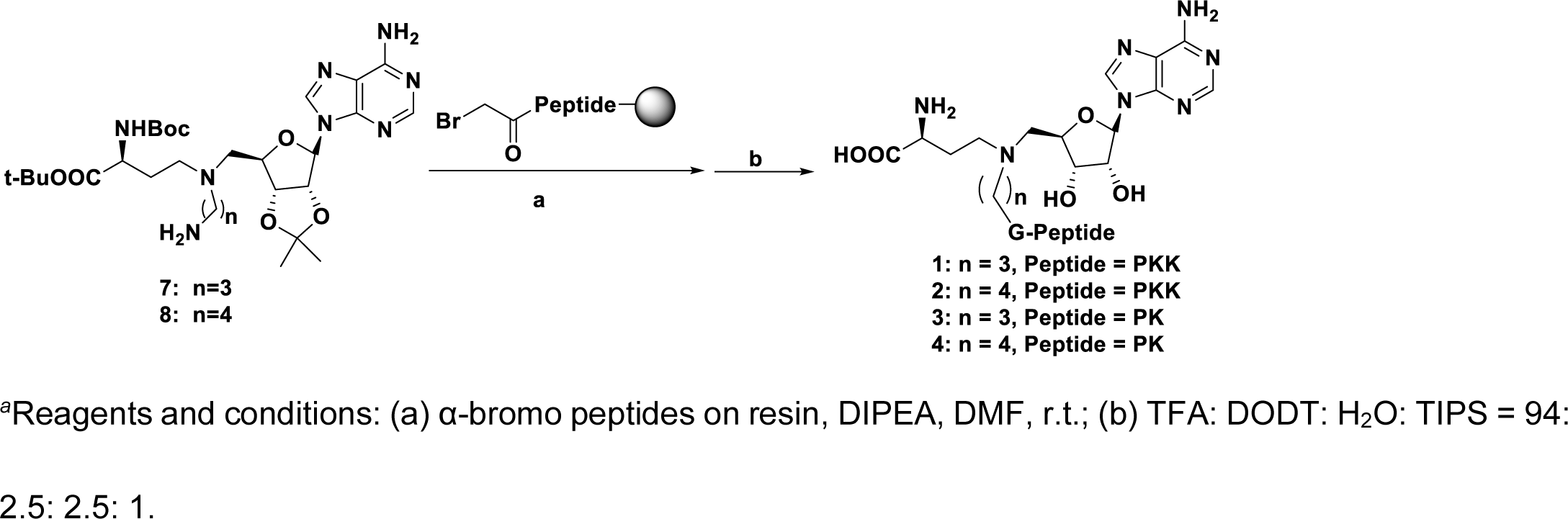
Synthetic route for compounds 1-4.*^a^*.

### Inhibition Evaluation

The inhibitory activities of compounds **1**-**4** against NTMT1 were evaluated in the SAH hydrolase (SAHH)-coupled fluorescence assay under the condition that both SAM and the peptide substrate GPKRIA were at their respective *K*_m_ values.^27^ Previous studies demonstrated that NAH-C4-GPKRIA and NAH-C4-GPKR bisubstrate analogs containing a 4-C carbon link showed slightly higher potency than those with a 3-C atom linker.^25^ However, the NAH-C3-GPKK bisubstrate analogue **1** with the 3-C atom linker (*K*_i, app_ = 7 ± 1 nM) exhibited similar inhibition as its 4-C linker analogue **2** (*K*_i, app_ = 5.0 ± 0.3 nM) (Table 1). Likewise, the NAH-C3-GPK bisubstrate analogue **3** with a 3-C carbon linker showed comparable inhibitory activity to compound **4** with a 4-C carbon linker, although the *K*_i, app_ values of **3** and **4** were increased by 6-8 fold compared to **1** and **2** due to the shortened peptide moiety from a tetrapeptide to a tripeptide.

**Table 1.** NTMT1 inhibition evaluation of bisubstrate analogues Inhibition Evaluation.

| Compound ID | Peptide | n | $K_{i, app}$ (nM) |
| --- | --- | --- | --- |
| **1 (NAH-C3-GPKK)** | GPKK | 3 | $7 \pm 1$ |
| **2 (NAH-C4-GPKK)** | GPKK | 4 | $5.0 \pm 0.3$ |
| **3 (NAH-C3-GPK)** | GPK | 3 | 41 ± 4 |
| **4 (NAH-C4-GPK)** | GPK | 4 | 44 ± 3 |

### MALDI-MS Methylation Inhibition Assay

For further validation, we evaluated the inhibitory effects of compound **1** on Nα-methylation progression with an orthogonal MALDI-MS methylation assay (Figure 2A-B).^24, 28^ As NTMT1 can trimethylate the Nα-amine group of the substrate peptide GPKRIA in which the Nα-amine group is a primary amine, there would be mo-, di-and tri-methylation states for the methylated product in the methylation assay. As the concentration of compound **1** increased, the tri-methylated GPKRIA decreased along with the increased level of unmethylated GPKRIA (Figure 2B). For compound **1**, tri-methylation of GPKRIA was almost abolished at 200 nM (Figure 2B). Also, all the methylation products of GPKRIA were reduced to less than 10% at 100 nM of compound **1** (Figure 2A-B). Through calculating the relative reaction rate based on the relative abundance with three species methylated products, quantitative analysis of the data and fitting them with different compound concentrations yielded the *K*_i, app_ of 15 ± 4 nM for **1** in the MS assay, which is comparable to the inhibitory activity in the fluorescence assay and further validated the potency of **1** for NTMT1.

**Figure 2.**
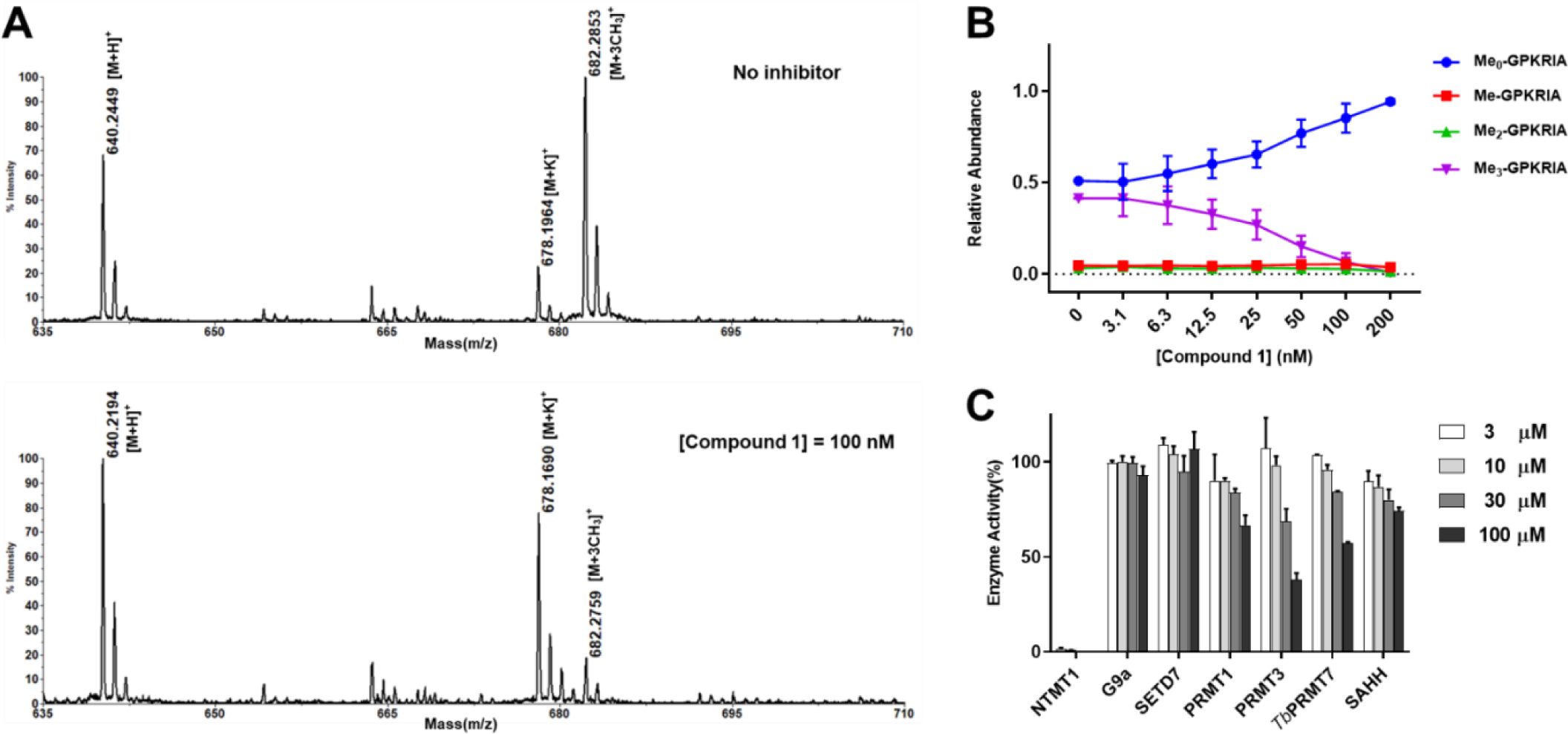
Methylation inhibition and selectivity study for **1**. (A) Representative MS results of MALDI-MS methylation inhibition assay for **1**; (B) Quantification of methylation progression of GPKRIA by NTMT1 in presence of various concentrations of **1** at 20 min (n=2). (C) Selectivity of **1** against several methyltransferases and SAH hydrolase (n=2).

### Selectivity Assessment

Next, we evaluated the selectivity of **1** for NTMT1 against a panel of several SAM-dependent methyltransferases, including two PKMTs (G9a and SETD7), and three PRMTs (PRMT1, PRMT3, and *Tb*PRMT7). We also include the SAHH, a coupling enzyme used in the fluorescence assay, which has a SAH binding site. Based on the results (Figure 2C), compound **1** retained the high selectivity to NTMT1, displaying more than 4,000-fold inhibition over other methyltransferases and SAHH.

### Co-crystal Structure of Compound 1 (NAH-C3-GPKK) with NTMT1

We obtained the X-ray crystal structure of the NTMT1-**1** complex (PDB ID: 6PVA) (Figure 3A), confirming its engagement with both cofactor and substrate binding sites of NTMT1. The structure of NTMT1-**1** overlaid well with the previously characterized co-crystal structure of bisubstrate inhibitor **NAH-C3-PPKRIA** (orange) with NTMT1 (PDB ID: 6DTN) (Figure 3B).^24^ The NAH moiety of **1** exhibited almost the same binding mode to NTMT1 as the NAH portion of **NAH-C3-PPKRIA** (Figure 3B), which is nearly identical with SAH in the ternary complex of substrate peptide/SAH with NTMT1 (PDB ID: 5E1M).^8^ Also, the peptide portion GPKK of **1** shows a similar binding mode compared with the PPKR peptide sequence in **NAH-C3-PPKRIA**. Specifically, the carbonyl oxygen of the first Gly forms a hydrogen bond with the side chain of Asn168 (Figure 3C). The second Pro residue inserts into the hydrophobic pocket formed by Leu31, Ile37, and Ile214, while the ε-amine group of the third Lys forms three hydrogen bonds with side chains of Asp177, Asp180, and Ser182.

**Figure 3.**
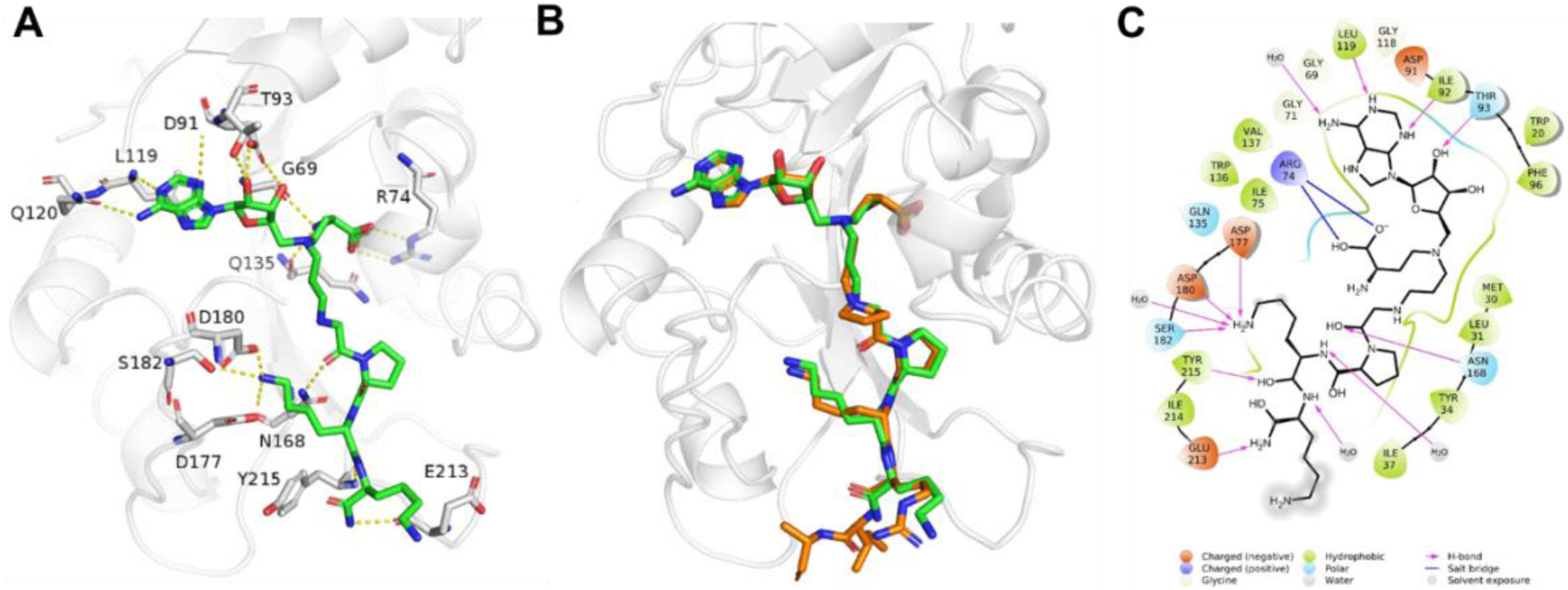
X-ray co-crystal structure of NTMT1-**1** (PDB ID: 6PVA). (A) Zoomed view of interactions of **1** (green) complex with NTMT1 (gray). H-bond interactions are shown in yellow dotted lines; (B) Structural alignment of NTMT1-**1** (gray/green) complex with NTMT1-**NAH-C3-PPKRIA** (orange) binary complex (PDB ID: 6DTN); (C) 2D interaction diagram (Schrödinger Maestro) of **1** with NTMT1.

### Design of Affinity Probe 6

As the last lysine is exposed to the solvent and its backbone forms interactions with NTMT1, we proposed the introduction of a clickable propargyl Gly (pG) to the C-terminus of the peptide of **1**, as well as linking the biotin group with pG to obtain the biotinylated probe, would retain the interaction with NTMT1 and allow the derivatization to immobilize the probe on streptavidin beads. Biotin-streptavidin is a powerful non-covalent interaction with a high binding affinity (*K*_d_ = ∼10^-14^ M).^29^ The biotin-streptavidin system has been widely applied in many biological applications, including proteome profiling, bio-imaging, biotinylated antibodies, and protein purification.^29^ Thus, probe **6** (NAH-C3-GPKK-biotin) was designed and synthesized for selectivity profiling (Scheme 2). To further confirm our design strategy, we also synthesized compound **5** (NAH-C3-GPKKpG) with a pG as the appendage (Scheme 2). Compound **5** demonstrated a comparable IC_50_ value (120 ± 20 nM) as compound **1** (150 ± 20 nM) in the SAHH-coupled fluorescence assay with a preincubation time of 10 minutes (Figure S1), supporting our design.

**Scheme 2.**
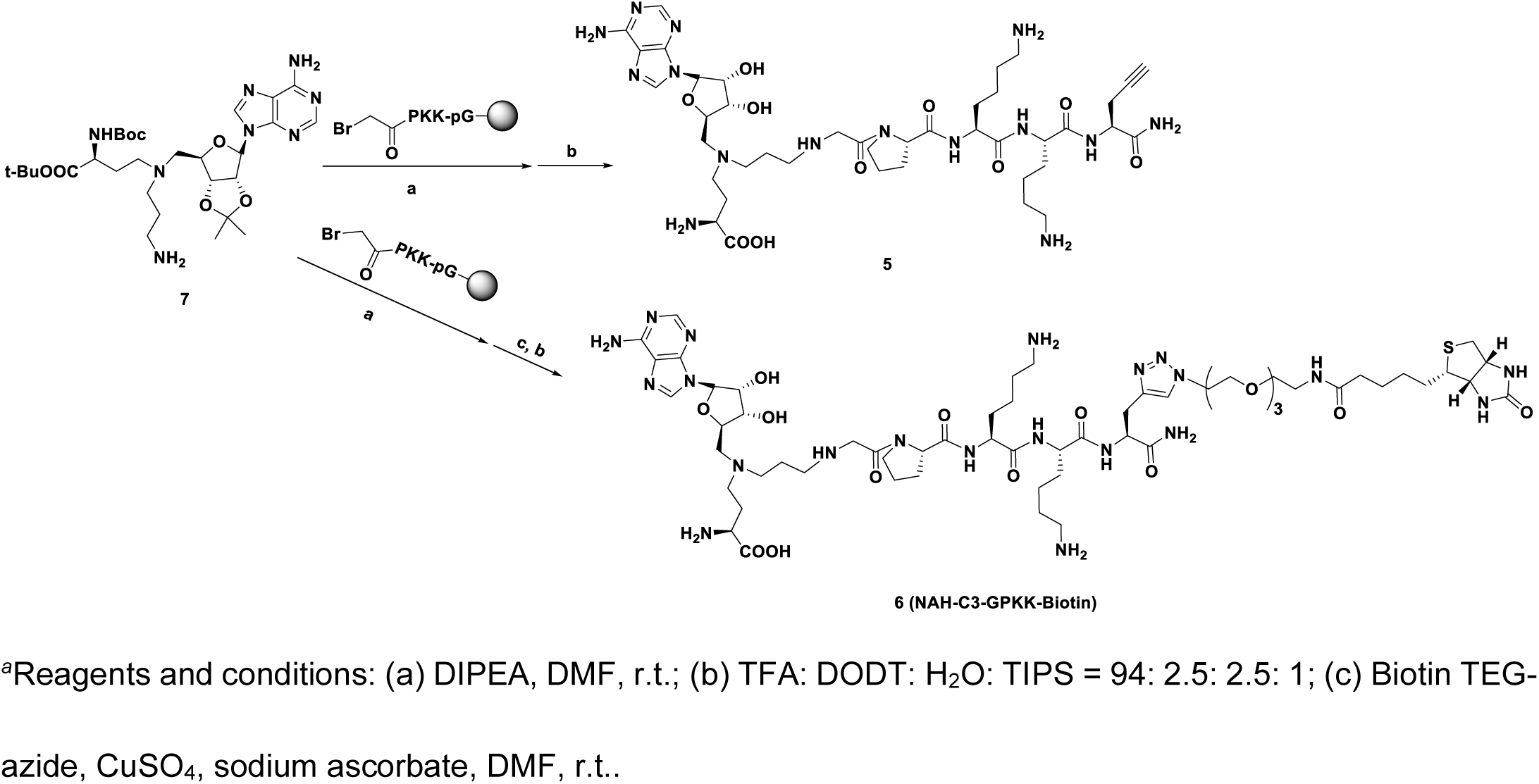
Synthetic route for compounds 5 and 6.*^a^*.

### Chemical Proteomic Profiling with Immobilized Probe 6

To investigate the optimal condition for the pulldown study, we first preincubated purified recombinant NTMT1 with immobilized probe **6** on streptavidin beads and eluted with a salt gradient in the elution buffer. NTMT1 was eluted off with the probe until the salt concentration reached 4 M NaCl, confirming strong interaction between **6** and NTMT1 (Figure S2). To remove non-specific and weak binders, we chose the lysis buffer containing 700 mM NaCl to wash the immobilized beads after preincubation with HeLa cell lysates. In parallel, the streptavidin beads alone were used as a negative control to remove any nonspecific binder. NTMT1 was enriched by probe **6** but not in the bead alone (Figure 4A). Specific enrichment of NTMT1 was further confirmed by competition experiments using the free inhibitor **1** (Figure 4A, Probe 6+1), indicating that pre-incubation of the lysate with **1** blocked the interaction between probe **6** and NTMT1 and allowed for the identification of specific interacting partners for probe **6**. NTMT1 was not washed off by the buffer containing 700 mM NaCl (Figure 4A, 700 mM wash lanes in probe **6**), consistent with the tight interaction between NTMT1 and probe **6**.

**Figure 4.**
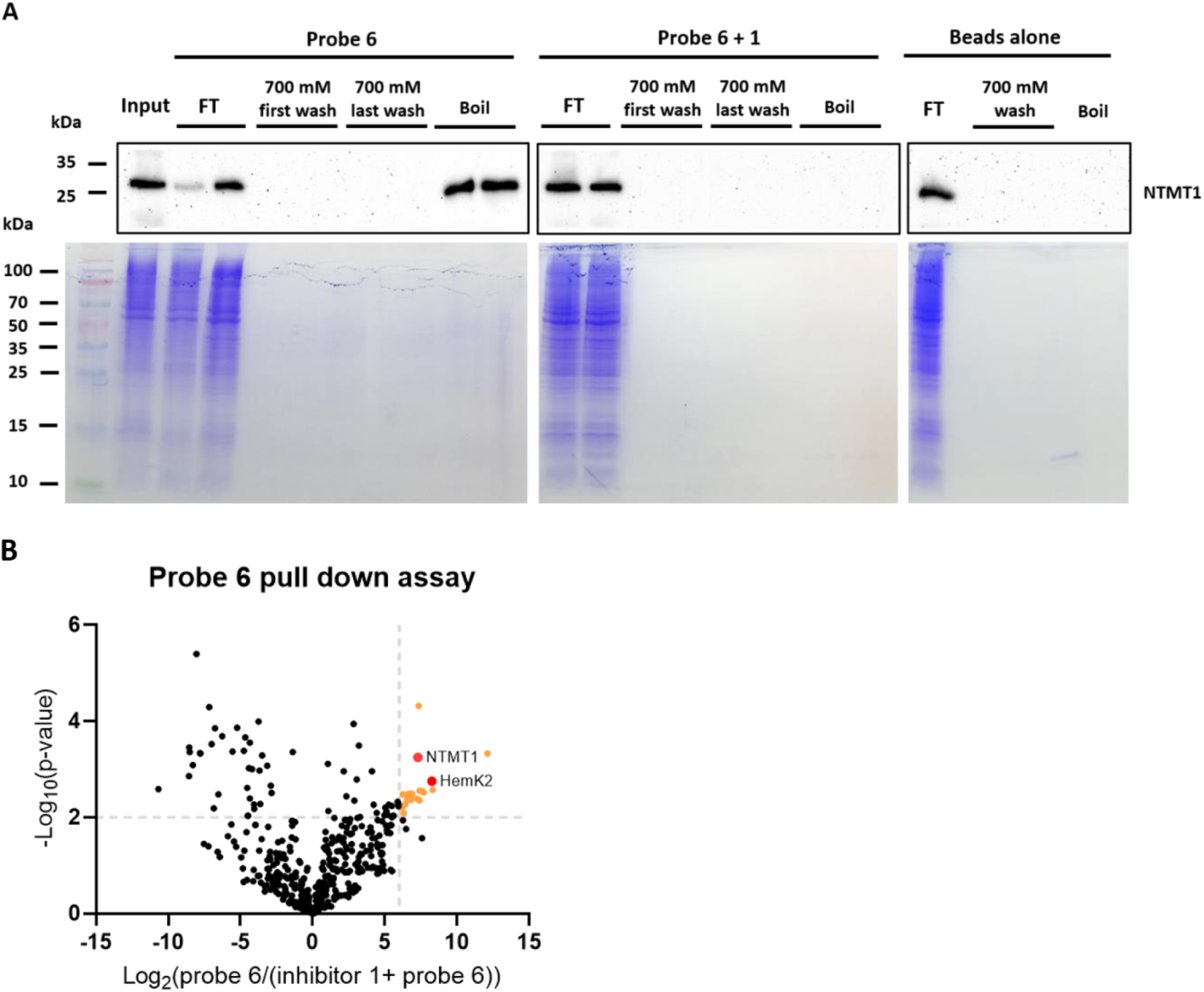
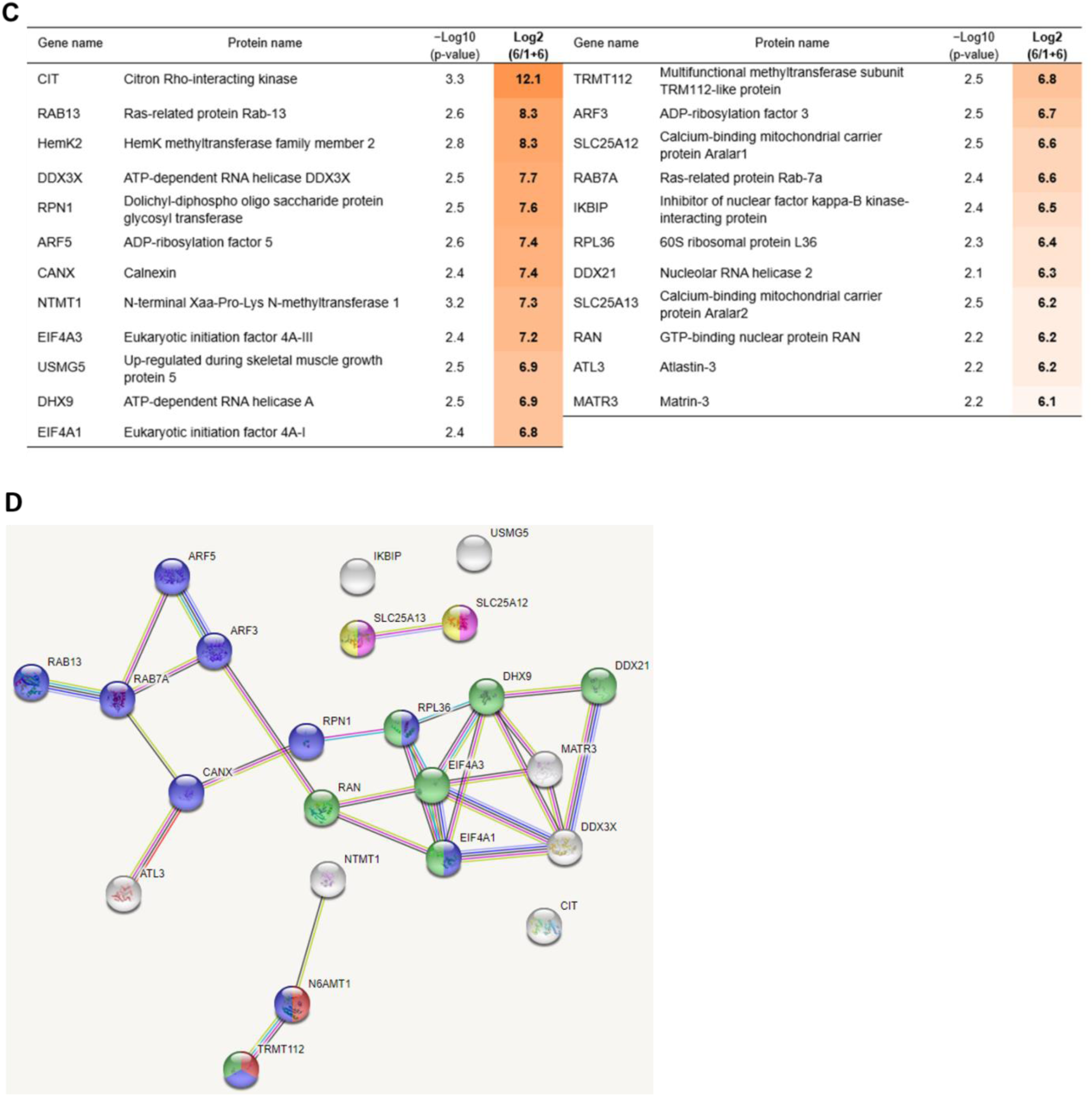
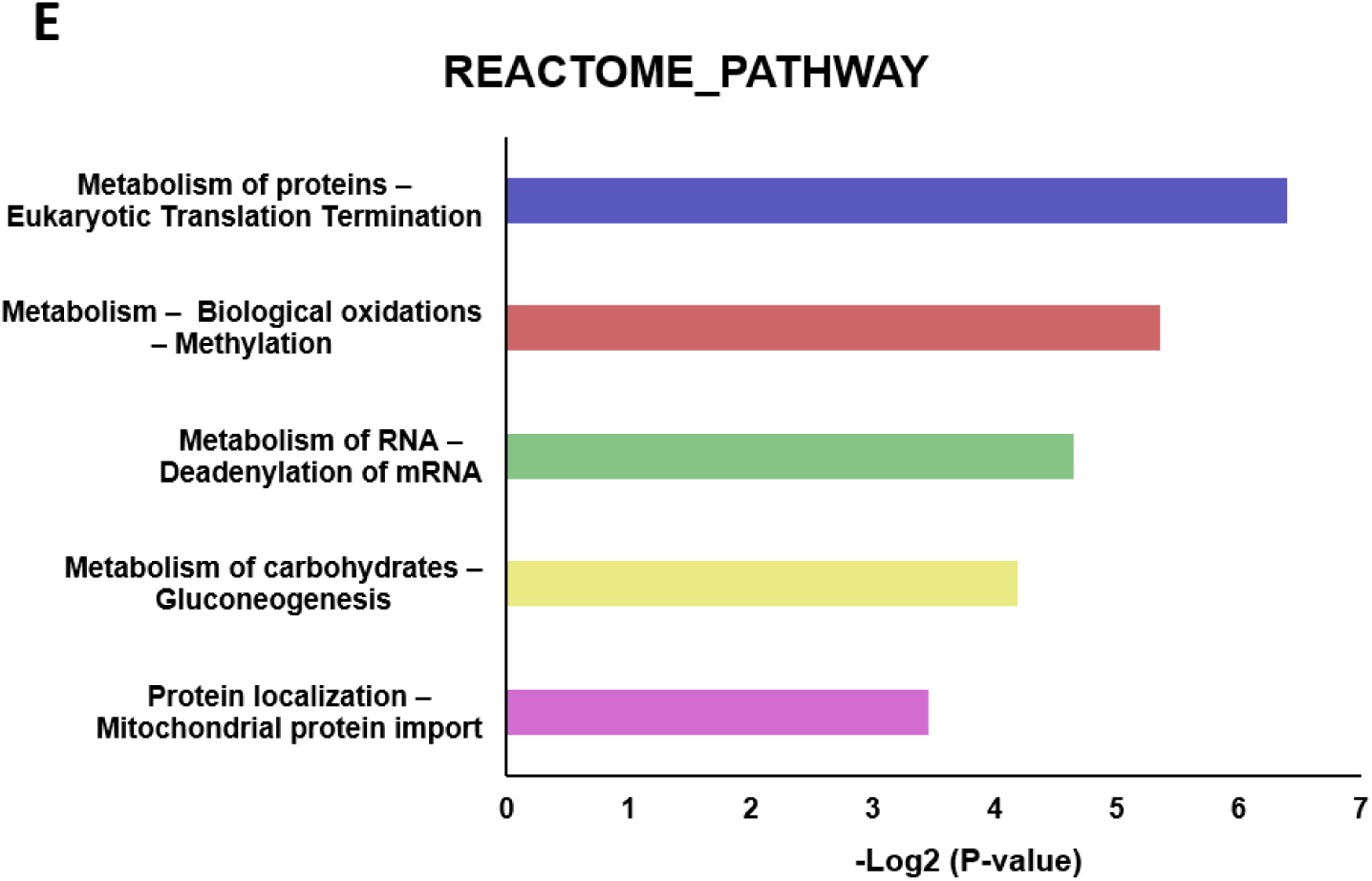
Competitive chemical proteomic profiling with immobilized probe **6**. (A) Probe **6** enriches NTMT1 from HeLa cell lysates. Western blot analysis of NTMT1 in HeLa cells lysates with the presence or absence of NTMT1 inhibitor **1** were incubated with immobilized probe **6**. The NTMT1 immunoblots of the flow-through (FT), 700 mM NaCl wash, and boiled beads with anti-NTMT1 antibody (Cell signaling #13432) were shown. (B) The volcano plot of −Log_10_(p-value) vs Log_2_(probe/competition) for proteins identified via LC-MS/MS using probe **6** and inhibitor **1**. Statistically significant interactors for probe **6** were defined as protein hits with fold change = Log_2_(Intensity (probe **6**)/Intensity (inhibitor **1** + probe **6**)) ≥ 6 and a confidence value = −log_10_(P-value) ≥ 2. Positions of NTMT1 and HemK2 are indicated in the plot by red circles. (C) The list of proteins that show statistically significant enrichment with probe **6** from HeLa cells. The third column shows confidence values (−log_10_(P-value)), and the fourth column shows enrichment scores. (D) STRING network of significant interactors for probe **6**. The STRING database was used to generate a network with these interactors. Each circle (node) represents a protein. Each line between the nodes represents a detected connection between two proteins in the STRING database. (E) Reactome pathway analysis of significant interactors for probe **6**. The interactors involved pathways were analyzed by STRING and DAVID databases. Proteins involve in each pathway were showed in the same color (Figure 4D) as the pathway bar. Proteins showed in white were not involved in any pathway from these two databases.

Having confirmed the specific enrichment of NTMT1 by immobilized probe **6**, we performed unbiased proteomic analysis of HeLa cell proteomics enriched by probe **6** through LC-MS/MS. We identified 23 proteins that were preferentially enriched by probe **6** with ≥6-fold change of (log_2_(probe 6/probe 6 + 1)) and ≥2 of confidence value (−log_10_(P-value)) for each identified protein (Figure 4B, Tables S1). Besides NTMT1, HemK methyltransferase family member 2 (HemK2, also known as KMT9^26, 30^ or N6AMT1^4, 30^), and its binding partner Trm112, are the only identified methyltransferase subunits among the 23 identified proteins (Figure 4B-C), supporting the relatively high selectivity of the NTMT1 bisubstrate inhibitor. Interestingly, both NTMT1 and HemK2 were identified as differentially expressed genes potentially involved in glioma stem cell DNA hypermethylation (Figure 4D).^31^ Other identified proteins included citron Rho-interacting kinase, small GTPases, RNA helicases, and ADP-ribosylation factors that have been involved in various pathways (Figure 4B-E). Notably, those identified proteins could either directly interact with the bait **6** or indirectly interact through NTMT1 or other possible targets. Validation on those interactions will be pursued through a pulldown study with purified NTMT1 or immunoprecipitation in the future. HemK2 and Trm112 forms a functional methyltransferase complex,^4, 26, 30^ with HemK2 as the catalytic subunit and Trm112 as an activator. The HemK2-Trm112 methyltransferase complex has a dual methyltransferase activity for either the protein lysine residue of histone H4 (named as KMT9) or glutamine residue of eRF1 (named as HemK2).^4, 26^ Although methyltransfer activity on a glutamine peptide (GGQSALR) is better than a lysine peptide (GKGGAKR) with both sequences conform to a similar motif as G-Q/K-XXX-K/R.^4, 26^

HemK2/KMT9 contains the SAM binding site and the substrate-binding site, while probe **6** consisted of a SAM analogue (NAH) and a peptide sequence GPKK containing two lysine residues. Thus, we hypothesized that the HemK2/KMT9 methyltransferase complex may bind with the NAH moiety or/and lysine-rich peptide sequence of the probe **6**. To validate this potential interaction, we tested the inhibitory activity of the parent compound **1** for the HemK2/KMT9 complex with a preferred GGQ substrate peptide.^30^ In Figure 5, compound **1** exhibited a potent inhibition for HemK2/KMT9 with an IC_50_ value of 0.30 ± 0.03 μM at 0.6 μM of the complex, which is tenfold more potent than a general methyltransferase inhibitor sinefungin (Figure 5A-B). When the concentration of the HemK2/KMT9 complex increased to 10 μM, compound **1** exhibited an IC_50_ value of 6.3 ± 0.8 μM, while a general methyltransferase inhibitor sinefungin only inhibited about 50% of activity at 100 μM (Figure 5C-D). As the IC_50_ value was approximately half of the enzyme concentration used in the assay in both cases, compound **1** acted as a tight inhibitor for the HemK2/KMT9 complex.^32^ To our knowledge, this is the first reported inhibitor for HemK2/KMT9 activity. Through monomethylation of histone H4 lysine 12 (H4K12), HemK2/KMT9 colocalized with H4K12me1 at gene promoters and regulated the gene transcription initiation.^26^ Overexpression of HemK2/KMT9/N6AMT1 was observed in prostate cancer cells, and knockdown of HemK2/KMT9/N6AMT1 strongly reduced the proliferation of prostate cancer cell lines.^26^ Besides, knockdown of HemK2/KMT9/N6AMT1 severely inhibited the growth of prostate tumor xenografts.^26^ Giving those emerging implications of HemK2/KMT9/N6AMT1 in prostate cancers, compound **1** would provide a rational lead for the development of inhibitors targeting the HemK2/KMT9 methyltransferase complex to treat protest cancer, as well as further improve the selectivity of NTMT1 inhibitors.

**Figure 5.**
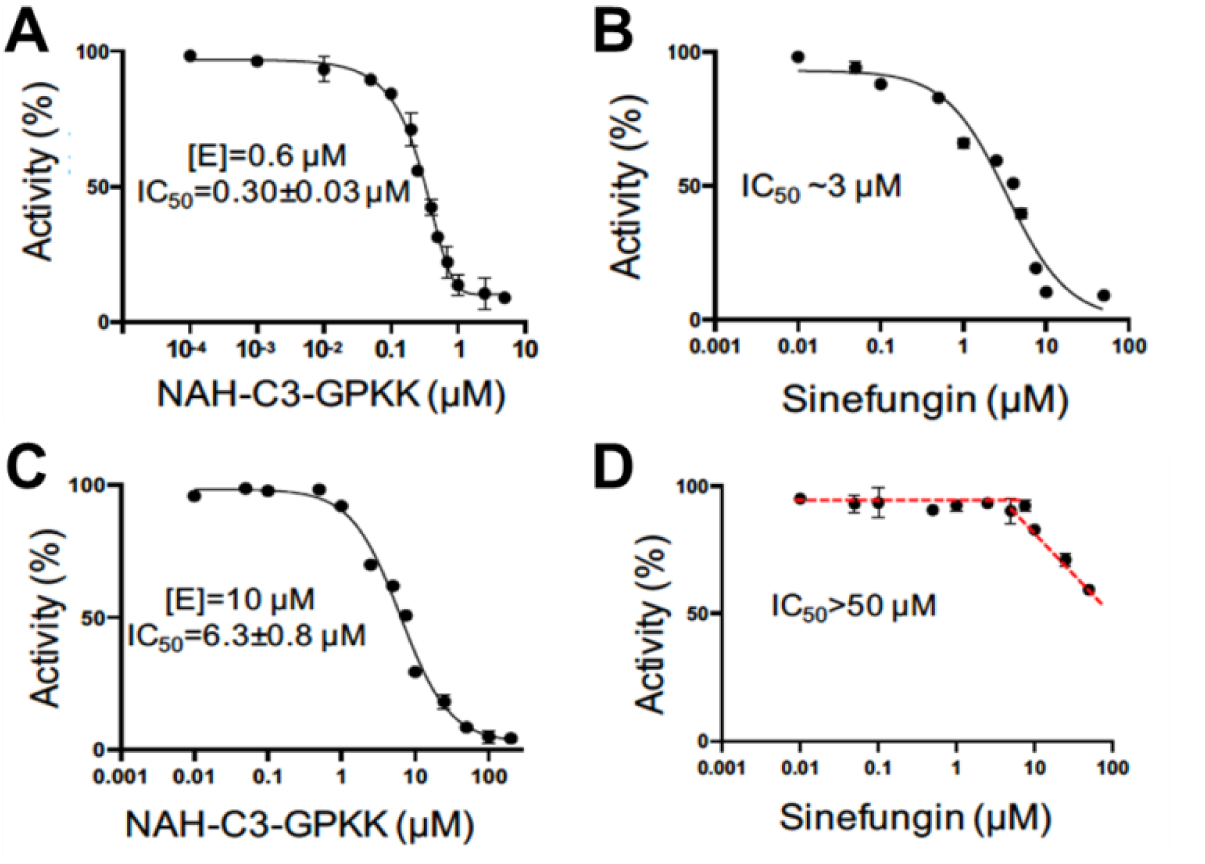
IC_50_ curves of compound **1** (NAH-C3-GPKK) and sinefungin for HemK2/KMT9-Trm112 activity on a GGQ peptide (n=3). (A) and (B) IC_50_ curves of compound **1** (NAH-C3-GPKK) and sinefungin at 0.6 μM of the complex; (C) and (D) IC_50_ curves of compound **1** (NAH-C3-GPKK) and sinefungin at 10 μM of the complex.

## CONCLUSION

In summary, we have rationally developed an affinity probe **6** based on the parent compound **1** that is a highly potent and relatively selective bisubstrate inhibitor for NTMT1. Chemproteomic studies with probe **6** confirmed the high selectivity of the NTMT1 bisubstrate inhibitor since the HemK2/KMT9-Trm112 methyltransferase complex was the only other methyltransferase that was enriched by probe **6**. Subsequent validation confirmed the parent inhibitor **1** was a tight inhibitor for the HemK2/KMT9 activity. As the first inhibitor for HemK2/KMT9, compound **1** would be a valuable probe and lead to developing potent inhibitors for HemK2/KMT9. Additionally, the future molecular mechanism investigation of **1** for HemK2/KMT9 would afford a solid basis to develop selective inhibitors for either methyltransferase.

## METHODS

### Chemistry General Procedures

The reagents and solvents were purchased from commercial sources (Fisher) and used directly. Final compounds were purified on preparative high-pressure liquid chromatography (RP-HPLC) was performed on Agilent 1260 Series system. Systems were run with 0-20% methanol/water gradient with 0.1% TFA. Matrix-assisted laser desorption ionization mass spectra (MALDI-MS) data were acquired in positive-ion mode using a Sciex 4800 MALDI TOF/TOF MS. The peptides were synthesized using a CEM Liberty Blue Automated Microwave Peptide Synthesizer with the manufacturer’s standard coupling cycles at 0.1 mmol scales. The purity of the final compounds was confirmed by Agilent 1260 Series HPLC system. Systems were run with a 0-40% methanol/water gradient with 0.1% TFA. All the purity of target compounds showed >95%.

*α-bromo peptides (G*PK, G*PKK, G*PKKpG).* To the solution of bromo acetic acid (139 mg, 1 mmol) in DMF(3 mL), DIC (155 μL, 1 mmol) was added and the solution was stirred for 20min at r.t. Then added the solution into the peptides on the resin (0.1 mmol). The mixture was shaken overnight at r.t. The suspension was filtered, and the resin was washed with DMF (3 ml x 3). Test cleavage (TFA: TIPS: H_2_O = 95:2.5:2.5) was performed to monitor the progress of the reaction.

NAH-C3-GPKK (**1**). To a suspension of G*PKK on the resin (0.1 mmol) in DMF (3 mL) was added compound **7** (91 mg, 0.15 mmol) and DIPEA (26 μL, 0.15 mmol). The mixture was shaken at room temperature overnight. The suspension was filtered, and the resin was subsequently washed with DMF (3 mL x 3), MeOH (3 mL x 3), and CH_2_Cl_2_ (3 mL x 3). The peptide conjugates were cleaved from the resin in the presence of a cleavage cocktail (10 mL) containing trifluoroacetic acid (TFA)/2,2’-(Ethylenedioxy)-diethanethiol (DODT) /triisopropylsilane (TIPS) /water (94:2.5:1:2.5 v/v) at room temperature for 4-5 h and resin was rinsed with a small amount of TFA. Volatiles were removed under N_2_ and the residue was precipitated with 10 vol. cold anhydrous ether, and collected by centrifugation. The supernatant was discarded and the pellet was washed with chilled ether and air-dried. The white precipitate was dissolved in ddH_2_O and purified by reverse phase HPLC using a waters system with an XBridge (BEH C18, 5 μm, 10 × 250 mm) column with 0.1% TFA in water (A) and MeOH (B) as the mobile phase with monitoring at 214 nm and 254 nm. MALDI-MS (positive) m/z: calcd for C_36_H_63_N_14_O_9_ [M + H]^+^ m/z 835.4902, found m/z 835.5424.

NAH-C4-GPKK (**2**). The compound was prepared according to the procedure for NAH-C3-GPKK (**1**) and purified by reverse-phase HPLC. MALDI-MS (positive) m/z: calcd for C_37_H_65_N_14_O_9_ [M + H]^+^ m/z 849.5059, found m/z 849.7359.

NAH-C3-GPK (**3**). The compound was prepared according to the procedure for NAH-C3-GPKK (**1**) and purified by reverse-phase HPLC. MALDI-MS (positive) m/z: calcd for C_30_H_51_N_12_O_8_ [M + H]^+^ m/z 707.3953, found m/z 707.5096.

NAH-C4-GPK (**4**). The compound was prepared according to the procedure for NAH-C3-GPKK (**1**) and purified by reverse-phase HPLC. MALDI-MS (positive) m/z: calcd for C_31_H_53_N_12_O_8_ [M + H]^+^ m/z 721.4109, found m/z 721.5273.

NAH-C3-GPKKpG (**5**). The compound was prepared according to the procedure for NAH-C3-GPKK (**1**) and purified by reverse-phase HPLC. MALDI-MS (positive) m/z: calcd for C_41_H_68_N_15_O_10_ [M + H]^+^ m/z 930.5274, found m/z 930.6367.

NAH-C3-GPKK-Biotin (**6**). To a suspension of G*PKK-pG on the resin (0.05 mmol) in DMF (2 mL) was added compound **7** (45 mg, 0.075 mmol) and DIPEA (13 μL, 0.075 mmol). The mixture was shaken at room temperature overnight. The suspension was filtered, and the resin was subsequently washed with DMF (3 mL x 3), MeOH (3 mL x 3), and CH_2_Cl_2_ (3 mL x 3). Then CuSO_4_·5H_2_O (25 mg, 0.1 mmol), sodium ascorbate (20 mg, 0.1mmol), and DMF (2 mL) were mixed with the resin. The mixture was shaken overnight at room temperature. The cleavage and purification of the conjugate were conducted according to the procedure for NAH-C3-GPKK (**1**). MALDI-MS (positive) m/z: calcd for C_59_H_100_N_21_O_15_S [M + H]^+^ m/z 1374.7428, found m/z 1375.1959.

### Protein Expression and Purification

Expression and purification of NTMT1, G9a, SETD7, PRMT1, PRMT3, *Tb*PRMT7, and HemK2-Trm112 were performed as previously described.^8, 25, 30, 33–35^

### NTMT1 Biochemical Assays

A fluorescence-based SAHH-coupled assay was applied to study the IC_50_ and *K*_i,app_ values of compounds with both SAM and the peptide substrate at their *K*_m_ values. For 10min pre-incubation, the methylation assay was performed under the following conditions in a final well volume of 40 μL: 25 mM Tris (pH = 7.5), 50 mM KCl, 0.01% Triton X-100, 5 μM SAHH, 0.1 μM NTMT1, 3 μM SAM, and 10 μM ThioGlo4. The inhibitors were added at concentrations ranging from 0.15 nM to 10 μM. After 10 min incubation, reactions were initiated by the addition of 0.5 μM GPKRIA peptide. For 120min pre-incubation, NTMT1 and compounds were pre-incubated for 120min. Then the pre-incubation mixture was added into reaction mixture: 25 mM Tris (pH = 7.5), 50 mM KCl, 0.01% Triton X-100, 5 μM SAHH, 3 μM SAM, and 10 μM ThioGlo4. The reactions were initiated by the addition of 0.5 μM GPKRIA peptide. Fluorescence was monitored on a BMG CLARIOstar microplate reader with excitation of 400 nm and emission of 465 nm. Data were processed by using GraphPad Prism software 7.0.

### Selectivity Assays

The selectivity studies of G9a, SETD7, PRMT1, PRMT3, *Tb*PRMT7, and SAHH were performed as previously described.^24, 25, 36^

### MALDI-MS Methylation Inhibition Assay

The methylation inhibition study of compound **1** with NTMT1 was performed as previously described.^25^

### HemK2 Methylation Inhibition Assay

The methylation inhibition assays of HemK2 was performed in a 20 μL reaction mixture containing 20 mM Tris-HCl (pH 8.0), 50 mM NaCl, 1 mM DTT and 20 μM SAM at 37°C, using high or low enzyme concentrations (10 μM or 0.6 μM), varied concentrations of eRF1 peptide (residues 179-183: KHGRGGQSALRFARL) (10 μM or 400 μM) and of compound 1 (NAH-C3-GPKK) or Sinefungin. The SAM-dependent methylation activity was measured by the Promega bioluminescence assay.^30^

### Pulldown Assay

Probe **6** (100 μL, 100 μM) was incubated with 50 μL streptavidin resin overnight in 20mM Tris, 50 mM KCl, pH=7.5. After washing out the nonbounded compound, purified NTMT1 was incubated with the probe-covered resin in 20 mM Tris, 50 mM KCl, 1 mM DTT, pH=7.5 for 4 h. Different concentration of the salt solution (500 mM, 1 M, 2 M, 4 M NaCl) was used to elute NTMT1.

HeLa cells (∼ 6×108 cells/mL) were lysed with fresh IP buffer containing 25 mM Tris, 150 mM NaCl, 1% NP-40, 1 mM EDTA, 5% glycerol, protease inhibitor (Halt ^TM^ protease inhibitor cocktail) pH 7.4. The cell suspension was incubated in rotation for 1 h at 4 °C and centrifuged at 15,000 g for 25 min at 4 °C to obtain the supernatant. The concentration of protein was determined with a BSA protein assay. The HeLa cell lysate was incubated with probe-covered resin overnight. For the competitor pulldown assay, competitor compound **1** (10 mM, 5 μL) was treated to the cell lysate first and incubate with compound-covered resin overnight. Resin alone was used as a negative control. Washed the resin five times with 1 mL IP buffer five times to remove the nonbounded proteins. The resin was then washed with 700 mM NaCl five times to remove the weak binding proteins. 1X SDS loading buffer was added and the resin was boiled for 5 min at 95 °C to elute proteins on it. Protein samples (10 μL) were loaded on 12.5 % SDS-PAGE gels and run at 120 V until the proteins come to the edge of the gel. Coomassie blue was used to stain the gels. Proteins on the gel were transferred to the PVDF membrane. After 5% milk blocking, the membrane was incubated with anti-NTMT1 (1:1000 dilution in 5% BSA in TBST buffer, Cell signaling #13432) at 4 °C overnight. Anti-rabbit IgG, HRP-linked antibody (1:1000 dilution in 5% BSA in TBST buffer, Cell signaling #7074) was incubated at 4 °C for 1h. The membrane was developed with ECL substrate (BIO-RAD #1705061) and imaged with a Proteinsimple FluorChem R.

Protein samples (10 μL) were loaded on 12.5 % SDS-PAGE gels and run at 120 V for 10 min. Protein bands were cut and submitted to the Laboratory for Biological Mass Spectrometry at Indiana University for LC-MS/MS.

### Co-crystallization and Structure Determination

The co-crystallization and structure determination of compound **1** with NTMT1 were performed as previously described.^24, 25^ Purified NTMT1 (30 mg/mL) was mixed with **1** at a 1:2 molar ratio, and the mixture was incubated overnight at 4°C. Using the hanging-drop vapor diffusion method and incubation at 20°C, the crystals of **1** were grown in 1.5 M lithium sulfate, 0.1 M sodium HEPES (pH = 7.5). The crystals were then harvested and flash frozen in liquid nitrogen. Data were collected on a single crystal at 12.0 keV at the GM/CA-CAT ID-D beamline at the Advanced Photon Source, Argonne National Laboratory. The data were processed (Table S2) and the structure was solved as previously described.^24, 25^

## ASSOCIATED CONTENT

### Supporting Information

MS and HPLC analysis of compounds **1**-**6**; Figure S1. IC_50_ curves for compounds **1** and **5**; Figure S2. Elution of recombinant NTMT1 from probe **6** coupled resin; Table S1. Proteomic analysis of HeLa cell proteomics enriched by probe **6** through LC-MS/MS; Table S2. Crystallography data and refinement statistics (PDB ID: 6PVA).

### Accession Codes

The coordinates for the structure of human NTMT1 in complex with compound **NAH-C3-GPKK** (PDB ID: 6PVA) has been deposited in the Protein Data Bank.

### Author Information

**Corresponding Author** *Phone: (765) 494 3426. E-mail

### Author Contributions

R.H. conceived and directed the project. R.H. and D.C. designed the compounds. D.C. synthesized, purified, and characterized the compounds. D.C. purified NTMT1 and performed co-crystallization. N.N. collected X-ray diffraction data and refined the co-crystal structure. Y.M. performed the pulldown study and bioinformatic analysis. X.C. supervised and D.Y. characterized the inhibition study for HemK2-Trm112. D.C. and Y.M. wrote the manuscript. R.H. and X.C. edited the manuscript. All authors provided critical feedback and have approved the final version of the manuscript.

### Notes

The authors declare no competing financial interest.

### Funding

The authors acknowledge the support from NIH grants R01GM117275 (RH), 1R01GM127896 (NN), 1R01AI127793 (NN), and R35GM134744 (XC). XC, a CPRIT Scholar in Cancer Research, thanks the support of Cancer Prevention and Research Institute of Texas (RR160029). GM/CA@APS beamline has been funded in whole or in part with Federal funds from the National Cancer Institute (ACB-12002) and the National Institute of General Medical Sciences (AGM-12006).

## ACKNOWLEDGMENT

GM/CA@APS has been funded by the National Cancer Institute (ACB-12002) and the National Institute of General Medical Sciences (AGM-12006, P30GM138396). This research used resources of the Advanced Photon Source, a U.S. Department of Energy (DOE) Office of Science User Facility operated for the DOE Office of Science by Argonne National Laboratory under Contract No. DE-AC02-06CH11357. We also thank supports from the Department of Medicinal Chemistry and Molecular Pharmacology (RH) and the Department of Biological Sciences (NN) at Purdue University. We thank Dr. Clayton B. Woodcock for providing purified human HemK2-Trm112 complex.

## ABBREVIATIONS

NTMT: protein N-terminal methyltransferase
SAM: *S*-adenosyl-L-methionine
SAH: S-adenosyl-L-homocysteines
SAHH: SAH hydrolase
MT: methyltransferase
PKMT: protein lysine methyltransferase;
PRMT: protein arginine methyltransferase
G9a: euchromatic histone-lysine N-methyltransferase 2
SETD7: SET domain-containing protein 7
*Tb*PRMT7: *Trypanosoma brucei* protein arginine methyltransferase 7
HemK2: HemK methyltransferase family member 2
rt: room temperature
TFA: trifluoroacetic acid.

## Supporting Information

### MS and HPLC analysis of compounds 1-6

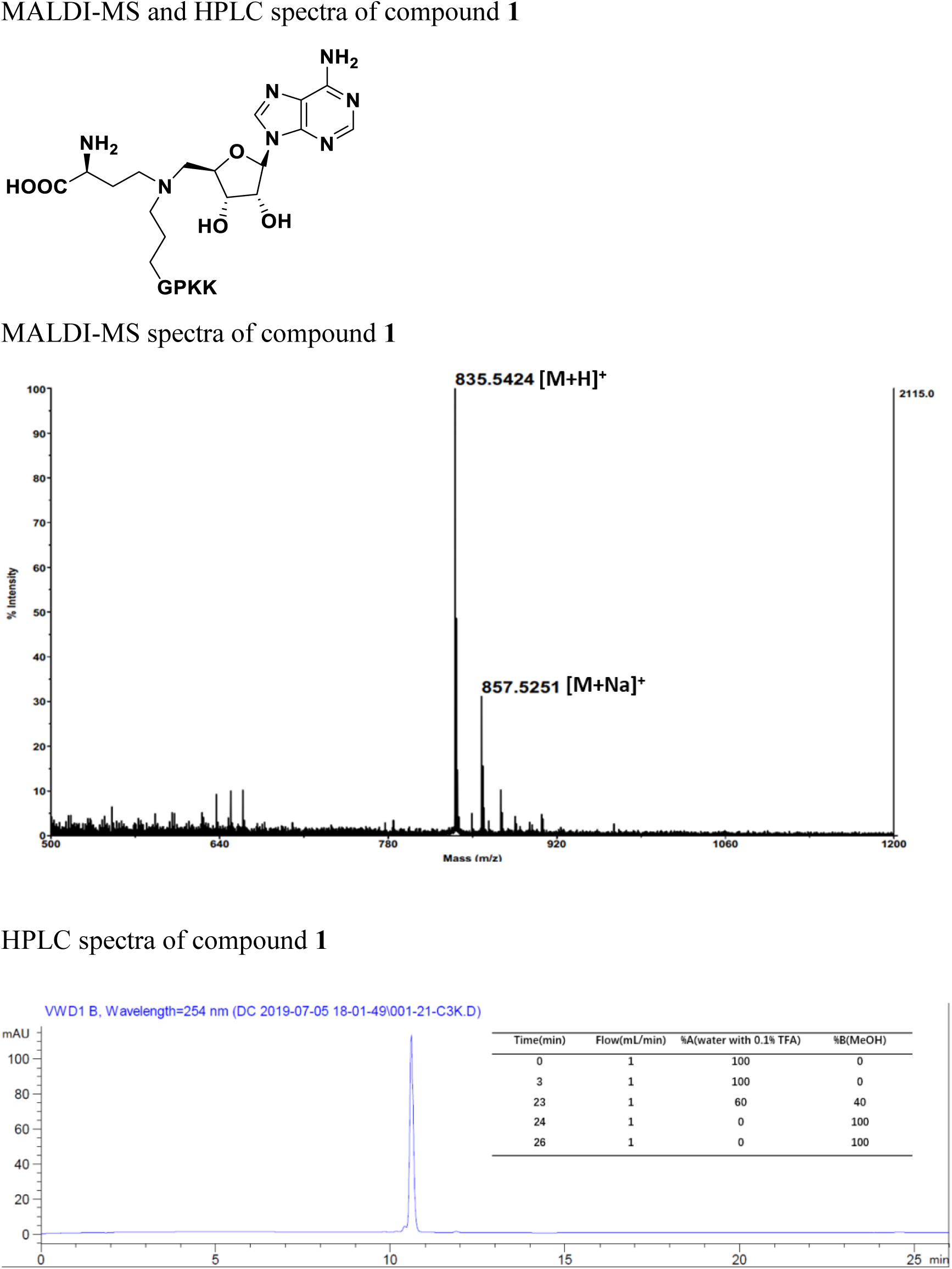
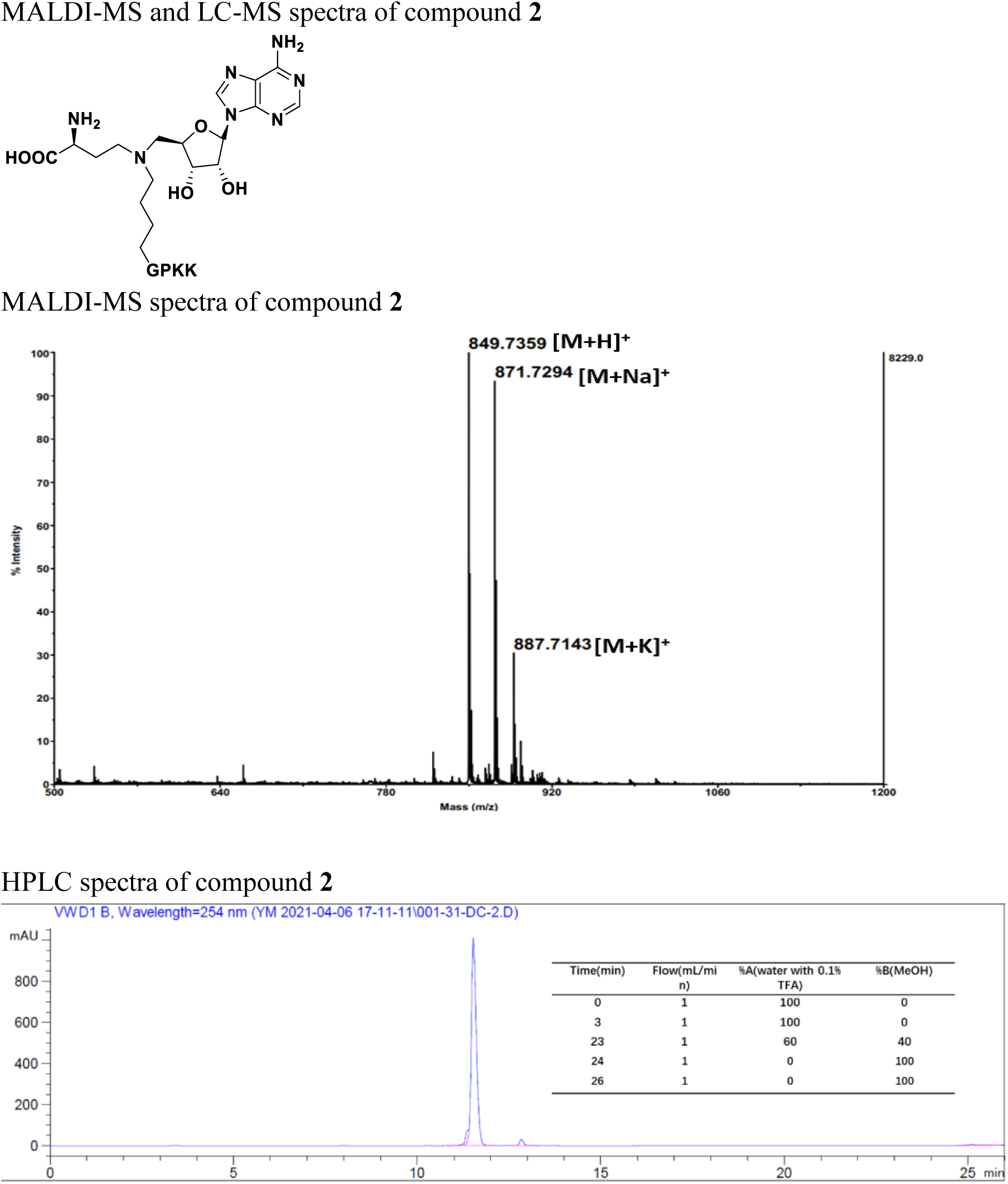
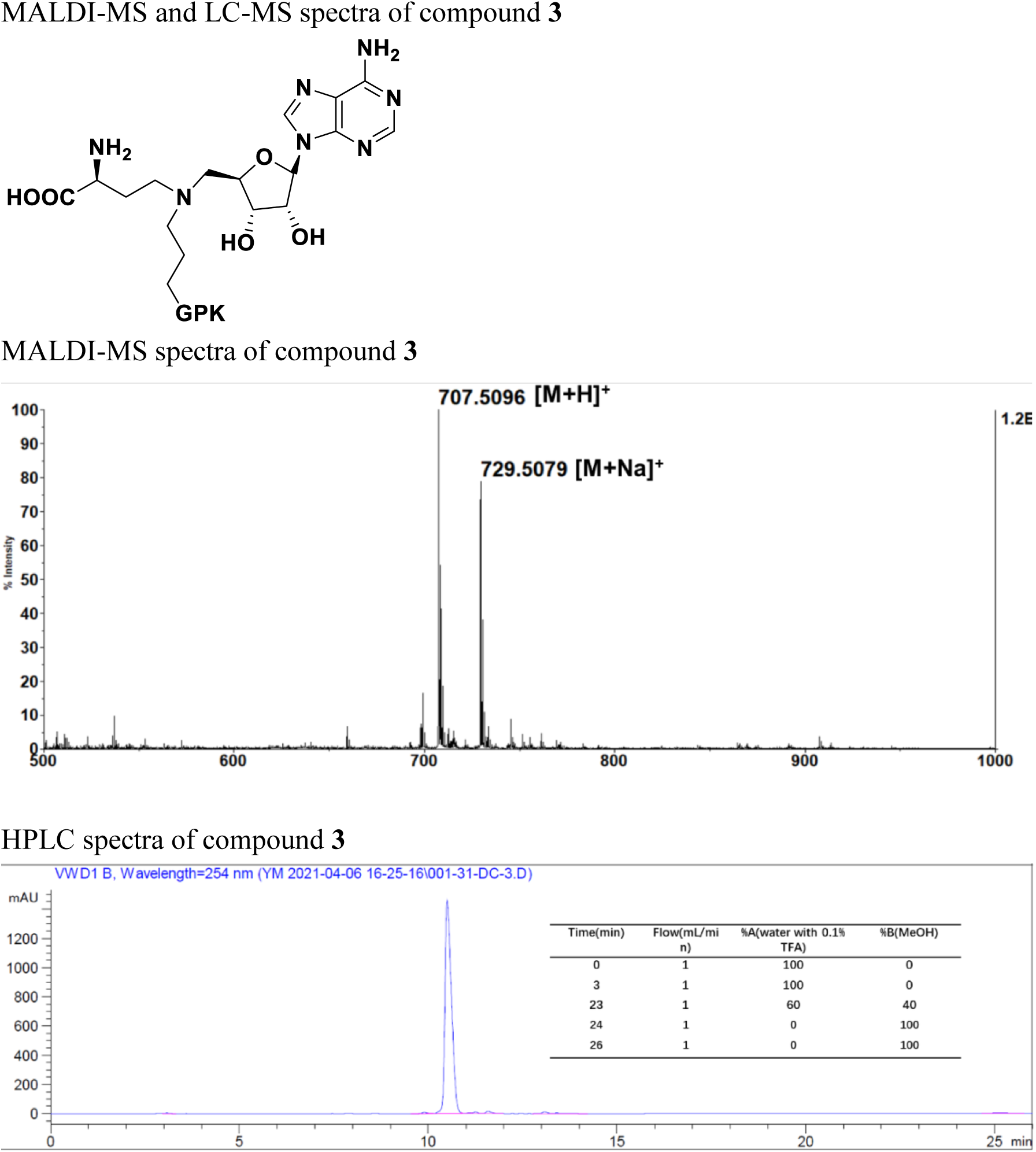
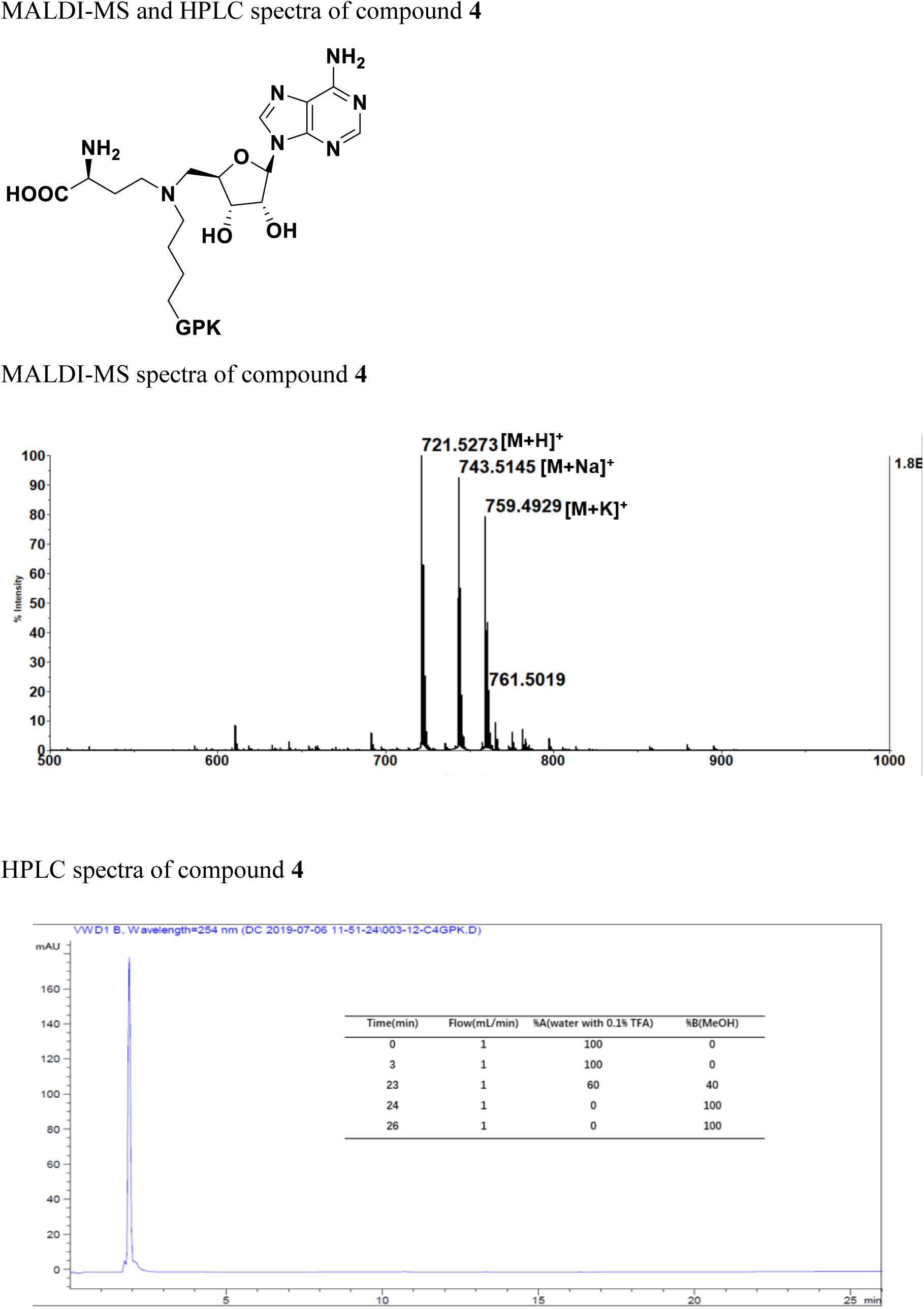
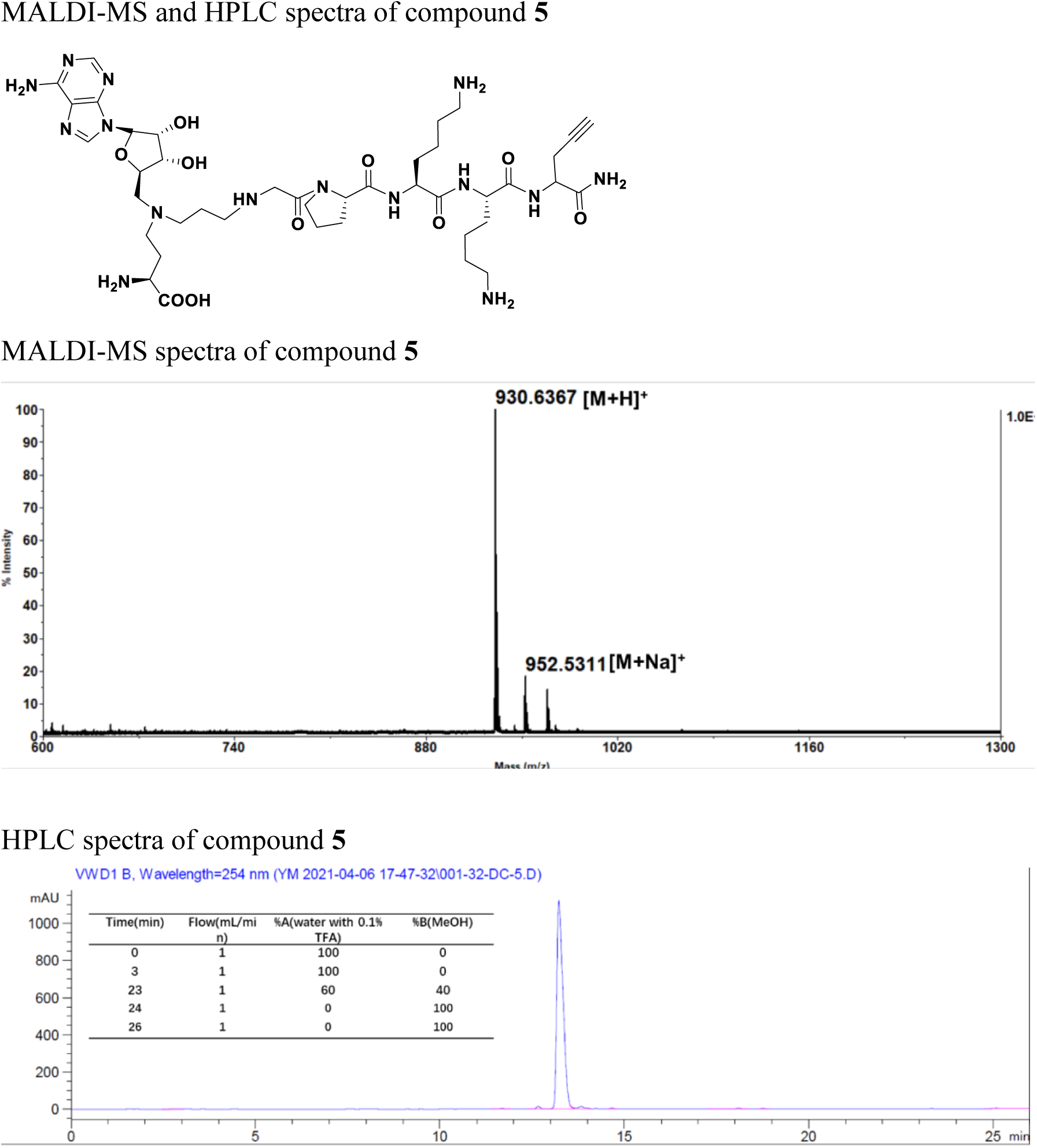
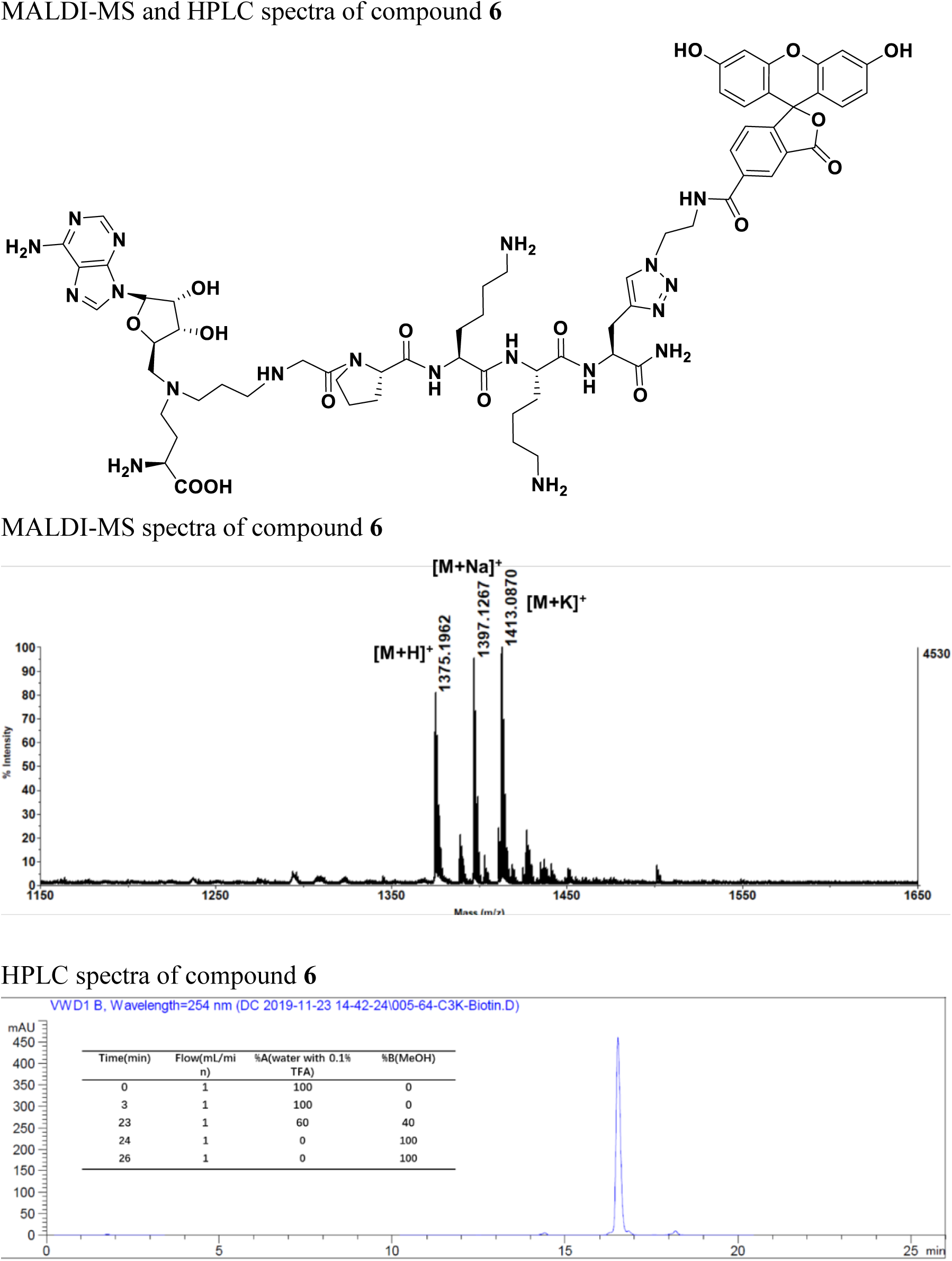

**Supporting Figure S1.**
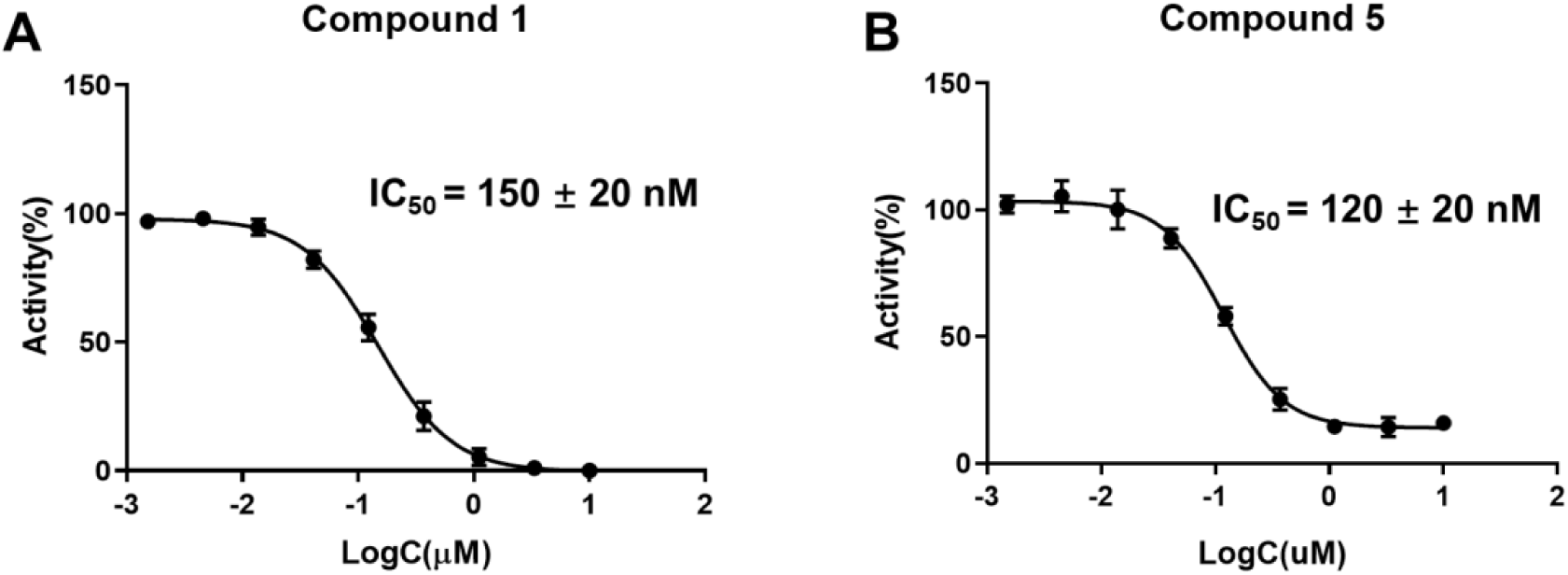
IC_50_ curves for compounds **1** (A) and **5** (B), respectively.

**Supporting Figure S2.**
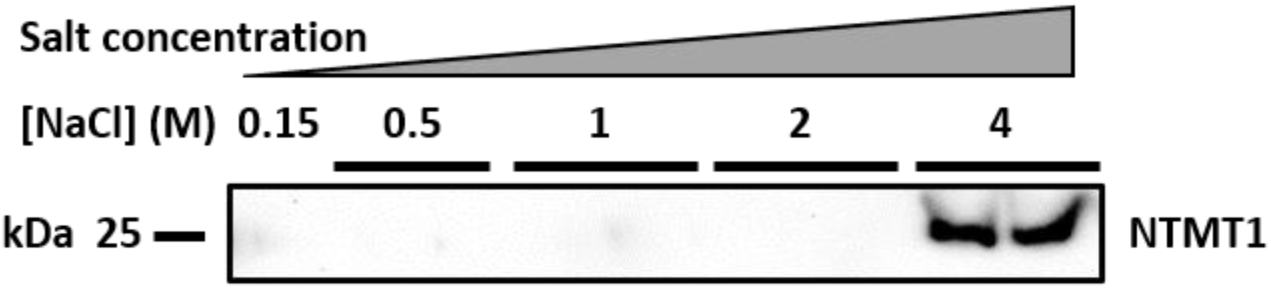
Elution of recombinant NTMT1 from probe **6** coupled resin. Probe 6 shows tight binding with NTMT1. Recombinant NTMT1 were incubated with probe 6 coupled streptavidin beads and eluted with different concentration of salt solution (0.15, 0.5, 1, 2, and 4 M).

**Supporting Table S1.** Proteomic analysis of HeLa cell proteomics enriched by probe 6 through LC-MS/MS.

| Accession | Protein name | Fold change = Log2(probe 6/ (inhibitor 1 + probe 6)) | Confidence value = $-\log_{10}(\text{P-value})$ |
| --- | --- | --- | --- |
| O14578-3 | Isoform 3 of Citron Rho-interacting kinase [OS=Homo sapiens] | 12.1 | 3.3 |
| Q9Y5N5 | HemK methyltransferase family member 2 [OS=Homo sapiens] | 8.3 | 2.8 |
| P51153 | Ras-related protein Rab-13 [OS=Homo sapiens] | 8.3 | 2.6 |
| O00571 | ATP-dependent RNA helicase DDX3X [OS=Homo sapiens] | 7.7 | 2.5 |
| Q96AG4 | Leucine-rich repeat-containing protein 59 [OS=Homo sapiens] | 7.6 | 1.6 |
| P04843 | Dolichyl-diphosphooligosaccharide--protein glycosyltransferase subunit 1 [OS=Homo sapiens] | 7.6 | 2.5 |
| P84085 | ADP-ribosylation factor 5 [OS=Homo sapiens] | 7.4 | 2.6 |
| P27824-2 | Isoform 2 of Calnexin [OS=Homo sapiens] | 7.4 | 2.4 |
| Q9BV86-1 | N-terminal Xaa-Pro-Lys N-methyltransferase 1 [OS=Homo sapiens] | 7.3 | 3.2 |
| P38919 | Eukaryotic initiation factor 4A-III [OS=Homo sapiens] | 7.2 | 2.4 |
| Q08211 | Atp-dependent rna helicase a [OS=Homo sapiens] | 6.9 | 2.5 |
| Q96IX5 | Up-regulated during skeletal muscle growth protein 5 [OS=Homo sapiens] | 6.9 | 2.5 |
| P60842 | Eukaryotic initiation factor 4A-I [OS=Homo sapiens] | 6.8 | 2.4 |
| Q9UI30 | Multifunctional methyltransferase subunit TRM112-like protein [OS=Homo sapiens] | 6.8 | 2.5 |
| P61204 | ADP-ribosylation factor 3 [OS=Homo sapiens] | 6.7 | 2.5 |
| P51149 | ras-related protein Rab-7a [OS=Homo sapiens] | 6.6 | 2.4 |
| **Q70UQ0-4** | **Isoform 4 of Inhibitor of nuclear factor kappa-B kinase-interacting protein [OS=Homo sapiens]** | **6.5** | **2.4** |
| **Q9Y3U8** | **60S ribosomal protein L36 [OS=Homo sapiens]** | **6.4** | **2.3** |
| **P13073** | **Cytochrome c oxidase subunit 4 isoform 1, mitochondrial [OS=Homo sapiens]** | **6.4** | **1.7** |
| **Q9NR30-1** | **Nucleolar RNA helicase 2 [OS=Homo sapiens]** | **6.3** | **2.1** |
| **P62826** | **GTP-binding nuclear protein RAN [OS=Homo sapiens]** | **6.2** | **2.2** |
| **P43243** | **Matrin-3 [OS=Homo sapiens]** | **6.2** | **2.2** |
| **O75746** | **Calcium-binding mitochondrial carrier protein Aralar1 [OS=Homo sapiens]** | **6.2** | **2.5** |
| **Q9UJS0-2** | **Isoform 2 of Calcium-binding mitochondrial carrier protein Aralar2 [OS=Homo sapiens]** | **6.2** | **2.5** |
| **P00403** | **Cytochrome c oxidase subunit 2 [OS=Homo sapiens]** | **6.2** | **1.9** |
| **Q6DD88** | **Atlastin-3 [OS=Homo sapiens]** | **6.1** | **2.3** |
| **P51148-2** | **Isoform 2 of Ras-related protein Rab-5C [OS=Homo sapiens]** | **6.0** | **2.3** |
| **Q96A26** | **Protein FAM162A [OS=Homo sapiens]** | **5.9** | **2.3** |
| **O00567** | **Nucleolar protein 56 [OS=Homo sapiens]** | **5.9** | **2.3** |
| **Q9NZ45** | **CDGSH iron-sulfur domain-containing protein 1 [OS=Homo sapiens]** | **5.7** | **2.0** |
| **P61224-1** | **Ras-related protein Rap-1b [OS=Homo sapiens]** | **5.6** | **2.1** |
| **Q14697-2** | **Isoform 2 of Neutral alpha-glucosidase AB [OS=Homo sapiens]** | **5.6** | **2.2** |
| **Q9Y277-2** | **Isoform 2 of Voltage-dependent anion-selective channel protein 3 [OS=Homo sapiens]** | **5.5** | **0.9** |
| **O75947-1** | **ATP synthase subunit d, mitochondrial [OS=Homo sapiens]** | **5.5** | **1.8** |
| **Q9BVP2** | **Guanine nucleotide-binding protein-like 3 [OS=Homo sapiens]** | **5.5** | **2.0** |
| **Q16790** | **Carbonic anhydrase 9 [OS=Homo sapiens]** | **5.4** | **1.8** |
| O60762 | Dolichol-phosphate<br>mannosyltransferase subunit 1<br>[OS=Homo sapiens] | 5.4 | 1.7 |
| O60506 | Heterogeneous nuclear<br>ribonucleoprotein Q [OS=Homo<br>sapiens] | 5.4 | 0.9 |
| O43169 | Cytochrome b5 type B [OS=Homo<br>sapiens] | 5.3 | 1.9 |
| Q9BWF3-1 | RNA-binding protein 4 [OS=Homo<br>sapiens] | 5.2 | 2.1 |
| Q9Y2X3 | Nucleolar protein 58 [OS=Homo<br>sapiens] | 5.2 | 2.1 |
| P36542-1 | ATP synthase subunit gamma,<br>mitochondrial [OS=Homo sapiens] | 5.2 | 1.7 |
| P33993-1 | DNA replication licensing factor MCM7<br>[OS=Homo sapiens] | 5.1 | 2.0 |
| O43809 | Cleavage and polyadenylation specificity<br>factor subunit 5 [OS=Homo sapiens] | 5.1 | 1.7 |
| P12956 | X-ray repair cross-complementing<br>protein 6 [OS=Homo sapiens] | 5.1 | 2.2 |
| Q07065 | Cytoskeleton-associated protein 4<br>[OS=Homo sapiens] | 5.0 | 2.1 |
| P01891 | HLA class I histocompatibility antigen,<br>A-68 alpha chain [OS=Homo sapiens] | 5.0 | 1.9 |
| P51648-2 | Isoform 2 of Fatty aldehyde<br>dehydrogenase [OS=Homo sapiens] | 5.0 | 2.2 |
| P22626 | heterogeneous nuclear<br>ribonucleoproteins A2/B1 [OS=Homo<br>sapiens] | 4.9 | 1.0 |
| P16615 | Sarcoplasmic/endoplasmic reticulum<br>calcium ATPase 2 [OS=Homo sapiens] | 4.9 | 0.9 |
| Q9NVI7-2 | Isoform 2 of ATPase family AAA<br>domain-containing protein 3A<br>[OS=Homo sapiens] | 4.9 | 1.4 |
| Q9Y4P3 | Transducin beta-like protein 2<br>[OS=Homo sapiens] | 4.9 | 2.0 |
| P20340-1 | Ras-related protein Rab-6A [OS=Homo<br>sapiens] | 4.8 | 1.2 |
| P61026 | ras-related protein rab-10 [OS=Homo<br>sapiens] | 4.7 | 1.3 |
| P11476-1 | Ras-related protein Rab-1A [OS=Homo sapiens] | 4.7 | 1.3 |
| P21796 | voltage-dependent anion-selective channel protein 1 [OS=Homo sapiens] | 4.7 | 1.1 |
| P61106 | Ras-related protein Rab-14 [OS=Homo sapiens] | 4.7 | 1.3 |
| O00264 | Membrane-associated progesterone receptor component 1 [OS=Homo sapiens] | 4.7 | 0.9 |
| P78527 | DNA-dependent protein kinase catalytic subunit [OS=Homo sapiens] | 4.6 | 0.9 |
| P34897-1 | Serine hydroxymethyltransferase, mitochondrial [OS=Homo sapiens] | 4.6 | 0.9 |
| P69905 | Hemoglobin subunit alpha [OS=Homo sapiens] | 4.6 | 1.6 |
| P63173 | 60s ribosomal protein l38 [OS=Homo sapiens] | 4.6 | 1.4 |
| P62491-1 | Ras-related protein Rab-11A [OS=Homo sapiens] | 4.5 | 1.9 |
| P30508 | HLA class I histocompatibility antigen, Cw-12 alpha chain [OS=Homo sapiens] | 4.5 | 1.4 |
| Q12905 | Interleukin enhancer-binding factor 2 [OS=Homo sapiens] | 4.5 | 0.9 |
| P62273-2 | Isoform 2 of 40S ribosomal protein S29 [OS=Homo sapiens] | 4.4 | 0.8 |
| Q8NBJ4 | Golgi membrane protein 1 [OS=Homo sapiens] | 4.4 | 1.9 |
| Q9BVK6 | Transmembrane emp24 domain-containing protein 9 [OS=Homo sapiens] | 4.4 | 1.6 |
| P16989-1 | Y-box-binding protein 3 [OS=Homo sapiens] | 4.3 | 1.8 |
| P05556-1 | Integrin beta-1 [OS=Homo sapiens] | 4.3 | 0.9 |
| P46940 | Ras GTPase-activating-like protein IQGAP1 [OS=Homo sapiens] | 4.2 | 2.3 |
| O95292 | Vesicle-associated membrane protein-associated protein B/C [OS=Homo sapiens] | 4.2 | 1.0 |
| P09669 | Cytochrome c oxidase subunit 6C [OS=Homo sapiens] | 4.2 | 0.9 |
| **Q9P0L0-2** | **Isoform 2 of Vesicle-associated membrane protein-associated protein A [OS=Homo sapiens]** | **4.2** | **1.5** |
| **O00461** | **Golgi integral membrane protein 4 [OS=Homo sapiens]** | **4.2** | **1.6** |
| **Q9NZ01-1** | **Very-long-chain enoyl-CoA reductase [OS=Homo sapiens]** | **4.1** | **0.9** |
| **P06576** | **ATP synthase subunit beta, mitochondrial [OS=Homo sapiens]** | **4.1** | **3.0** |
| **P61978-2** | **Isoform 2 of Heterogeneous nuclear ribonucleoprotein K [OS=Homo sapiens]** | **4.0** | **1.2** |
| **P22087** | **rRNA 2'-O-methyltransferase fibrillarin [OS=Homo sapiens]** | **4.0** | **1.3** |
| **P62910** | **60S ribosomal protein L32 [OS=Homo sapiens]** | **3.9** | **1.6** |
| **Q9NXW2-2** | **Isoform 2 of DnaJ homolog subfamily B member 12 [OS=Homo sapiens]** | **3.9** | **1.8** |
| **O43390-2** | **Isoform 2 of Heterogeneous nuclear ribonucleoprotein R [OS=Homo sapiens]** | **3.8** | **1.0** |
| **P27635** | **60S ribosomal protein L10 [OS=Homo sapiens]** | **3.7** | **1.3** |
| **Q8NE86** | **Calcium uniporter protein, mitochondrial [OS=Homo sapiens]** | **3.6** | **1.0** |
| **O75367-1** | **Core histone macro-H2A.1 [OS=Homo sapiens]** | **3.5** | **1.0** |
| **P62805** | **histone H4 [OS=Homo sapiens]** | **3.5** | **1.4** |
| **Q9Y5S2** | **Serine/threonine-protein kinase MRCK beta [OS=Homo sapiens]** | **3.4** | **1.8** |
| **P01889** | **HLA class I histocompatibility antigen, B-7 alpha chain [OS=Homo sapiens]** | **3.3** | **0.9** |
| **P62753** | **40S RIBOSOMAL PROTEIN S6 [OS=Homo sapiens]** | **3.3** | **1.2** |
| **Q14254** | **Flotillin-2 [OS=Homo sapiens]** | **3.3** | **1.3** |
| **P42167** | **Lamina-associated polypeptide 2, isoforms beta/gamma [OS=Homo sapiens]** | **3.3** | **2.0** |
| **P17301** | **Integrin alpha-2 [OS=Homo sapiens]** | **3.3** | **1.0** |
| **Q96CW1** | **AP-2 complex subunit mu [OS=Homo sapiens]** | **3.2** | **1.2** |
| **P42677** | **40S ribosomal protein S27 [OS=Homo sapiens]** | **3.2** | **3.5** |
| **Q96CS3** | **FAS-associated factor 2 [OS=Homo sapiens]** | **3.2** | **0.9** |
| **Q9HDC9** | **Adipocyte plasma membrane-associated protein [OS=Homo sapiens]** | **3.1** | **0.8** |
| **P49748-3** | **Isoform 3 of Very long-chain specific acyl-CoA dehydrogenase, mitochondrial [OS=Homo sapiens]** | **3.1** | **1.1** |
| **P50914** | **60S ribosomal protein L14 [OS=Homo sapiens]** | **3.1** | **2.0** |
| **P27348** | **14-3-3 protein theta [OS=Homo sapiens]** | **3.1** | **0.5** |
| **Q9Y265** | **RuvB-like 1 [OS=Homo sapiens]** | **3.1** | **1.0** |
| **P62258-1** | **14-3-3 protein epsilon [OS=Homo sapiens]** | **3.1** | **0.5** |
| **P17844** | **probable ATP-dependent RNA helicase DDX5 [OS=Homo sapiens]** | **3.1** | **2.8** |
| **P61981** | **14-3-3 protein gamma [OS=Homo sapiens]** | **3.0** | **0.5** |
| **P19388** | **DNA-directed RNA polymerases I, II, and III subunit RPABC1 [OS=Homo sapiens]** | **3.0** | **0.6** |
| **O14929** | **histone acetyltransferase type B catalytic subunit [OS=Homo sapiens]** | **3.0** | **0.5** |
| **P60866-2** | **Isoform 2 of 40S ribosomal protein S20 [OS=Homo sapiens]** | **3.0** | **1.0** |
| **Q92841** | **Probable ATP-dependent RNA helicase DDX17 [OS=Homo sapiens]** | **2.9** | **1.8** |
| **Q71UM5** | **40S ribosomal protein S27-like [OS=Homo sapiens]** | **2.9** | **2.3** |
| **P31947-1** | **14-3-3 protein sigma [OS=Homo sapiens]** | **2.9** | **0.4** |
| **P14625** | **Endoplasmic reticulum chaperone [OS=Homo sapiens]** | **2.8** | **3.9** |
| **P02788** | **Lactotransferrin [OS=Homo sapiens]** | **2.7** | **0.5** |
| **P05023** | **Sodium/potassium-transporting ATPase subunit alpha-1 [OS=Homo sapiens]** | **2.7** | **1.7** |
| **Q9Y5M8** | **signal recognition particle receptor subunit beta [OS=Homo sapiens]** | **2.7** | **1.3** |
| P45880-1 | Isoform 1 of Voltage-dependent anion-selective channel protein 2 [OS=Homo sapiens] | 2.7 | 1.4 |
| P53680 | AP-2 complex subunit sigma [OS=Homo sapiens] | 2.7 | 0.5 |
| O75396 | Vesicle-trafficking protein SEC22b [OS=Homo sapiens] | 2.6 | 1.6 |
| P48047 | ATP synthase subunit O, mitochondrial [OS=Homo sapiens] | 2.6 | 2.0 |
| P63104-1 | 14-3-3 protein zeta/delta [OS=Homo sapiens] | 2.6 | 0.5 |
| P29401-2 | Isoform 2 of Transketolase [OS=Homo sapiens] | 2.6 | 0.9 |
| P14868 | Aspartate--tRNA ligase, cytoplasmic [OS=Homo sapiens] | 2.5 | 1.9 |
| O00425 | Insulin-like growth factor 2 mRNA-binding protein 3 [OS=Homo sapiens] | 2.5 | 0.6 |
| P31946 | 14-3-3 protein beta/alpha [OS=Homo sapiens] | 2.5 | 0.4 |
| P08237-3 | Isoform 3 of ATP-dependent 6-phosphofructokinase, muscle type [OS=Homo sapiens] | 2.4 | 0.3 |
| Q9Y6M9 | NADH dehydrogenase [ubiquinone] 1 beta subcomplex subunit 9 [OS=Homo sapiens] | 2.4 | 0.8 |
| P25705-1 | ATP synthase subunit alpha, mitochondrial [OS=Homo sapiens] | 2.4 | 1.7 |
| P39023 | 60S ribosomal protein L3 [OS=Homo sapiens] | 2.4 | 1.3 |
| Q9HA64 | Ketosamine-3-kinase [OS=Homo sapiens] | 2.4 | 1.8 |
| P32969 | 60S ribosomal protein L9 [OS=Homo sapiens] | 2.3 | 2.4 |
| Q8N1F7-1 | Nuclear pore complex protein Nup93 [OS=Homo sapiens] | 2.3 | 1.4 |
| P62424 | 60S ribosomal protein L7a [OS=Homo sapiens] | 2.3 | 1.2 |
| Q96A08 | Histone H2B type 1-A [OS=Homo sapiens] | 2.3 | 1.0 |
| **Q16891-2** | **Isoform 2 of MICOS complex subunit MIC60 [OS=Homo sapiens]** | **2.3** | **1.9** |
| **Q16891-4** | **Isoform 4 of MICOS complex subunit MIC60 [OS=Homo sapiens]** | **2.3** | **1.9** |
| **Q96S19-4** | **Isoform 4 of UPF0585 protein C16orf13 [OS=Homo sapiens]** | **2.2** | **0.9** |
| **Q9Y3D9** | **28S ribosomal protein S23, mitochondrial [OS=Homo sapiens]** | **2.2** | **1.1** |
| **P39656** | **Dolichyl-diphosphooligosaccharide--protein glycosyltransferase 48 kDa subunit [OS=Homo sapiens]** | **2.2** | **0.4** |
| **Q07020** | **60S ribosomal protein L18 [OS=Homo sapiens]** | **2.2** | **3.0** |
| **P83731** | **60S ribosomal protein L24 [OS=Homo sapiens]** | **2.1** | **1.3** |
| **P40939** | **Trifunctional enzyme subunit alpha, mitochondrial [OS=Homo sapiens]** | **2.1** | **1.5** |
| **P52597** | **Heterogeneous nuclear ribonucleoprotein F [OS=Homo sapiens]** | **2.1** | **1.6** |
| **P42765** | **3-ketoacyl-CoA thiolase, mitochondrial [OS=Homo sapiens]** | **2.0** | **0.4** |
| **P31689-1** | **DnaJ homolog subfamily A member 1 [OS=Homo sapiens]** | **2.0** | **1.8** |
| **P24539** | **ATP synthase F(0) complex subunit B1, mitochondrial [OS=Homo sapiens]** | **1.9** | **1.5** |
| **P23396-1** | **40S ribosomal protein S3 [OS=Homo sapiens]** | **1.9** | **1.3** |
| **P18085** | **ADP-ribosylation factor 4 [OS=Homo sapiens]** | **1.8** | **1.6** |
| **Q12797** | **Aspartyl/Asparaginyl beta-hydroxylase [OS=Homo sapiens]** | **1.8** | **0.4** |
| **Q86UE4** | **protein LYRIC [OS=Homo sapiens]** | **1.7** | **0.9** |
| **P62241** | **40S ribosomal protein S8 [OS=Homo sapiens]** | **1.7** | **1.3** |
| **Q9Y241** | **HIG1 domain family member 1A, mitochondrial [OS=Homo sapiens]** | **1.7** | **0.5** |
| **P50454** | **Serpin H1 [OS=Homo sapiens]** | **1.7** | **0.9** |
| **P18124** | **60S ribosomal protein L7 [OS=Homo sapiens]** | **1.6** | **1.1** |
| P08574 | Cytochrome c1, heme protein, mitochondrial [OS=Homo sapiens] | 1.6 | 0.4 |
| P10809 | 60 kDa heat shock protein, mitochondrial [OS=Homo sapiens] | 1.6 | 1.5 |
| P07195 | L-lactate dehydrogenase B chain [OS=Homo sapiens] | 1.6 | 0.3 |
| Q13492-1 | Phosphatidylinositol-binding clathrin assembly protein [OS=Homo sapiens] | 1.6 | 0.3 |
| Q92769 | Histone deacetylase 2 [OS=Homo sapiens] | 1.5 | 0.3 |
| P46781 | 40S ribosomal protein S9 [OS=Homo sapiens] | 1.5 | 1.1 |
| P04075 | fructose-bisphosphate aldolase A [OS=Homo sapiens] | 1.5 | 0.3 |
| Q9Y230 | RuvB-like 2 [OS=Homo sapiens] | 1.5 | 2.0 |
| P35268 | 60S ribosomal protein L22 [OS=Homo sapiens] | 1.5 | 0.9 |
| P08195-4 | Isoform 4 of 4F2 cell-surface antigen heavy chain [OS=Homo sapiens] | 1.5 | 0.7 |
| P61353 | 60S ribosomal protein L27 [OS=Homo sapiens] | 1.4 | 0.7 |
| Q15233 | Non-POU domain-containing octamer-binding protein [OS=Homo sapiens] | 1.4 | 0.6 |
| Q12931 | heat shock protein 75 kDa, mitochondrial [OS=Homo sapiens] | 1.4 | 0.2 |
| Q15365 | Poly(RC)-binding protein 1 [OS=Homo sapiens] | 1.4 | 0.9 |
| P62244 | 40S ribosomal protein S15a [OS=Homo sapiens] | 1.4 | 1.6 |
| P02786 | Transferrin receptor protein 1 [OS=Homo sapiens] | 1.3 | 1.5 |
| Q7Z2W4 | zinc finger CCCH-type antiviral protein 1 [OS=Homo sapiens] | 1.2 | 0.7 |
| Q14684 | Ribosomal RNA processing protein 1 homolog B [OS=Homo sapiens] | 1.2 | 1.0 |
| O95782 | AP-2 complex subunit alpha-1 [OS=Homo sapiens] | 1.2 | 0.6 |
| Q99567 | Nuclear pore complex protein Nup88 [OS=Homo sapiens] | 1.2 | 0.2 |
| **P46776** | **60S ribosomal protein L27a [OS=Homo sapiens]** | **1.2** | **1.0** |
| **P57721-1** | **Poly(rC)-binding protein 3 [OS=Homo sapiens]** | **1.2** | **1.1** |
| **Q00839** | **Heterogeneous nuclear ribonucleoprotein U [OS=Homo sapiens]** | **1.1** | **1.2** |
| **P31943** | **Heterogeneous nuclear ribonucleoprotein H [OS=Homo sapiens]** | **1.1** | **0.8** |
| **P63010-2** | **Isoform 2 of AP-2 complex subunit beta [OS=Homo sapiens]** | **1.1** | **0.8** |
| **P08238** | **Heat shock protein HSP 90-beta [OS=Homo sapiens]** | **1.1** | **2.1** |
| **P67809** | **Nuclease-sensitive element-binding protein 1 [OS=Homo sapiens]** | **1.1** | **0.2** |
| **P54136-1** | **arginine--tRNA ligase, cytoplasmic [OS=Homo sapiens]** | **1.1** | **1.3** |
| **O43175** | **D-3-phosphoglycerate dehydrogenase [OS=Homo sapiens]** | **1.1** | **3.1** |
| **Q9NX63** | **MICOS complex subunit MIC19 [OS=Homo sapiens]** | **1.0** | **1.4** |
| **P35232** | **Prohibitin [OS=Homo sapiens]** | **0.9** | **0.5** |
| **P62829** | **60S ribosomal protein L23 [OS=Homo sapiens]** | **0.9** | **0.4** |
| **Q01813** | **ATP-dependent 6-phosphofructokinase, platelet type [OS=Homo sapiens]** | **0.9** | **1.3** |
| **P05388** | **60S acidic ribosomal protein P0 [OS=Homo sapiens]** | **0.9** | **1.5** |
| **P52272** | **Heterogeneous nuclear ribonucleoprotein M [OS=Homo sapiens]** | **0.9** | **0.5** |
| **Q9UHX1-1** | **poly(U)-binding-splicing factor PUF60 [OS=Homo sapiens]** | **0.9** | **0.1** |
| **P18077** | **60S ribosomal protein L35a [OS=Homo sapiens]** | **0.9** | **1.7** |
| **P39019** | **40S ribosomal protein S19 [OS=Homo sapiens]** | **0.9** | **0.8** |
| **P28066-1** | **Proteasome subunit alpha type-5 [OS=Homo sapiens]** | **0.9** | **0.1** |
| **Q5QNW6-2** | **Isoform 2 of Histone H2B type 2-F**<br>**[OS=Homo sapiens]** | **0.8** | **0.5** |
| **P62277** | **40S ribosomal protein S13 [OS=Homo sapiens]** | **0.8** | **0.7** |
| **P62701** | **40S ribosomal protein S4, X isoform**<br>**[OS=Homo sapiens]** | **0.8** | **1.4** |
| **Q14980-1** | **nuclear mitotic apparatus protein 1**<br>**[OS=Homo sapiens]** | **0.8** | **0.4** |
| **Q09666-1** | **Neuroblast differentiation-associated**<br>**protein AHNAK [OS=Homo sapiens]** | **0.8** | **0.9** |
| **P46783** | **40S ribosomal protein S10 [OS=Homo sapiens]** | **0.7** | **1.1** |
| **Q99714-1** | **3-hydroxyacyl-CoA dehydrogenase**<br>**type-2 [OS=Homo sapiens]** | **0.7** | **0.9** |
| **P06899** | **Histone H2B type 1-J [OS=Homo sapiens]** | **0.7** | **0.4** |
| **P04406-1** | **glyceraldehyde-3-phosphate**<br>**dehydrogenase [OS=Homo sapiens]** | **0.7** | **0.2** |
| **P11940-1** | **Polyadenylate-binding protein 1**<br>**[OS=Homo sapiens]** | **0.7** | **1.0** |
| **P46778** | **60S ribosomal protein L21 [OS=Homo sapiens]** | **0.6** | **0.6** |
| **P62847-4** | **Isoform 4 of 40S ribosomal protein S24**<br>**[OS=Homo sapiens]** | **0.6** | **0.3** |
| **P62266** | **40S ribosomal protein S23 [OS=Homo sapiens]** | **0.6** | **0.6** |
| **Q99623** | **Prohibitin-2 [OS=Homo sapiens]** | **0.5** | **0.4** |
| **P18621-3** | **Isoform 3 of 60S ribosomal protein L17**<br>**[OS=Homo sapiens]** | **0.5** | **0.3** |
| **P68366** | **Tubulin alpha-4A chain [OS=Homo sapiens]** | **0.5** | **0.3** |
| **P68104** | **Elongation factor 1-alpha 1 [OS=Homo sapiens]** | **0.5** | **0.4** |
| **Q96PK6-1** | **RNA-binding protein 14 [OS=Homo sapiens]** | **0.5** | **0.5** |
| **P35613** | **Basigin [OS=Homo sapiens]** | **0.5** | **0.7** |
| **Q13573** | **SNW domain-containing protein 1**<br>**[OS=Homo sapiens]** | **0.4** | **0.1** |
| **P26641** | **elongation factor 1-gamma [OS=Homo sapiens]** | **0.4** | **0.6** |
| P84098 | 60S ribosomal protein L19 [OS=Homo sapiens] | 0.4 | 0.2 |
| P04083 | annexin A1 [OS=Homo sapiens] | 0.4 | 0.1 |
| Q9BWM7 | Sideroflexin-3 [OS=Homo sapiens] | 0.4 | 0.1 |
| P06493 | Cyclin-dependent kinase 1 [OS=Homo sapiens] | 0.3 | 0.3 |
| P06733-1 | alpha-enolase [OS=Homo sapiens] | 0.3 | 0.0 |
| P60900 | Proteasome subunit alpha type-6 [OS=Homo sapiens] | 0.3 | 0.1 |
| Q13310-3 | Isoform 3 of Polyadenylate-binding protein 4 [OS=Homo sapiens] | 0.3 | 0.5 |
| P62888 | 60S ribosomal protein L30 [OS=Homo sapiens] | 0.2 | 0.1 |
| P26373-1 | 60S ribosomal protein L13 [OS=Homo sapiens] | 0.2 | 0.3 |
| P07900-2 | Isoform 2 of Heat shock protein HSP 90-alpha [OS=Homo sapiens] | 0.2 | 0.4 |
| P62913 | 60S ribosomal protein L11 [OS=Homo sapiens] | 0.2 | 0.6 |
| P62750 | 60S ribosomal protein L23a [OS=Homo sapiens] | 0.2 | 0.0 |
| Q9BUF5 | Tubulin beta-6 chain [OS=Homo sapiens] | 0.2 | 0.4 |
| P04908 | histone H2A type 1-B/E [OS=Homo sapiens] | 0.2 | 0.1 |
| O43795 | Unconventional myosin-Ib [OS=Homo sapiens] | 0.2 | 0.1 |
| P27105 | erythrocyte band 7 integral membrane protein [OS=Homo sapiens] | 0.2 | 0.1 |
| Q12965 | unconventional myosin-Ie [OS=Homo sapiens] | 0.1 | 0.1 |
| P22695 | Cytochrome b-c1 complex subunit 2, mitochondrial [OS=Homo sapiens] | 0.1 | 0.0 |
| P83881 | 60S ribosomal protein L36a [OS=Homo sapiens] | 0.1 | 0.0 |
| Q13509 | tubulin beta-3 chain [OS=Homo sapiens] | 0.1 | 0.1 |
| Q92614-1 | Unconventional myosin-XVIIIa [OS=Homo sapiens] | 0.1 | 0.0 |
| P00338-3 | Isoform 3 of L-lactate dehydrogenase A chain [OS=Homo sapiens] | 0.0 | 0.0 |
| **P07910-1** | **Heterogeneous nuclear ribonucleoproteins C1/C2 [OS=Homo sapiens]** | **0.0** | **0.0** |
| **P14618** | **Pyruvate kinase PKM [OS=Homo sapiens]** | **0.0** | **0.0** |
| **Q9UKS6** | **Protein kinase C and casein kinase substrate in neurons protein 3 [OS=Homo sapiens]** | **0.0** | **0.0** |
| **P68371** | **Tubulin beta-4B chain [OS=Homo sapiens]** | **0.0** | **0.0** |
| **P36578** | **60S ribosomal protein L4 [OS=Homo sapiens]** | **0.0** | **0.0** |
| **P11021** | **78 kDa glucose-regulated protein [OS=Homo sapiens]** | **0.0** | **0.0** |
| **P19338** | **Nucleolin [OS=Homo sapiens]** | **-0.1** | **0.0** |
| **P05387** | **60S acidic ribosomal protein P2 [OS=Homo sapiens]** | **-0.1** | **0.1** |
| **P61254** | **60S ribosomal protein L26 [OS=Homo sapiens]** | **-0.2** | **0.1** |
| **Q01105** | **Protein SET [OS=Homo sapiens]** | **-0.2** | **0.0** |
| **P21333** | **Filamin-A [OS=Homo sapiens]** | **-0.2** | **0.1** |
| **P07814** | **Bifunctional glutamate/proline--tRNA ligase [OS=Homo sapiens]** | **-0.2** | **0.1** |
| **Q02878** | **60S ribosomal protein L6 [OS=Homo sapiens]** | **-0.2** | **0.1** |
| **Q14244-7** | **Isoform 7 of Ensconsin [OS=Homo sapiens]** | **-0.2** | **0.3** |
| **P0DMV8** | **heat shock 70 kDa protein 1A [OS=Homo sapiens]** | **-0.3** | **0.3** |
| **P36957** | **Dihydrolipoyllysine-residue succinyltransferase component of 2-oxoglutarate dehydrogenase complex, mitochondrial [OS=Homo sapiens]** | **-0.3** | **0.1** |
| **P62851** | **40S ribosomal protein S25 [OS=Homo sapiens]** | **-0.3** | **0.2** |
| **P61513** | **60S ribosomal protein L37a [OS=Homo sapiens]** | **-0.3** | **0.0** |
| **P62906** | **60S ribosomal protein L10A [OS=Homo sapiens]** | **-0.3** | **0.5** |
| P63096-1 | Guanine nucleotide-binding protein G(i) subunit alpha-1 [OS=Homo sapiens] | -0.3 | 0.3 |
| P05141 | ADP/ATP translocase 2 [OS=Homo sapiens] | -0.3 | 0.2 |
| P62249 | 40S ribosomal protein S16 [OS=Homo sapiens] | -0.3 | 0.3 |
| P07437 | tubulin beta chain [OS=Homo sapiens] | -0.3 | 0.4 |
| Q15046 | Lysine--tRNA ligase [OS=Homo sapiens] | -0.4 | 0.9 |
| P38646 | Stress-70 protein, mitochondrial [OS=Homo sapiens] | -0.4 | 0.9 |
| P35579-1 | Myosin-9 [OS=Homo sapiens] | -0.4 | 0.2 |
| P49411 | elongation factor Tu, mitochondrial [OS=Homo sapiens] | -0.5 | 0.5 |
| Q02543 | 60S ribosomal protein L18a [OS=Homo sapiens] | -0.5 | 0.2 |
| Q8WV41 | Sorting nexin-33 [OS=Homo sapiens] | -0.5 | 0.1 |
| P0C0S5 | Histone H2A.Z [OS=Homo sapiens] | -0.5 | 0.2 |
| Q13155 | aminoacyl tRNA synthase complex-interacting multifunctional protein 2 [OS=Homo sapiens] | -0.5 | 0.1 |
| Q69YQ0-1 | Cytospin-A [OS=Homo sapiens] | -0.5 | 0.3 |
| Q9BQE3 | Tubulin alpha-1C chain [OS=Homo sapiens] | -0.6 | 0.5 |
| Q4KMQ1 | Taperin [OS=Homo sapiens] | -0.6 | 0.2 |
| P06702 | Protein S100-A9 [OS=Homo sapiens] | -0.6 | 0.1 |
| Q9BVA1 | Tubulin beta-2B chain [OS=Homo sapiens] | -0.6 | 0.7 |
| P11142-1 | Heat shock cognate 71 kDa protein [OS=Homo sapiens] | -0.6 | 0.7 |
| P09651-1 | Heterogeneous nuclear ribonucleoprotein A1 [OS=Homo sapiens] | -0.6 | 0.4 |
| P55084 | Trifunctional enzyme subunit beta, mitochondrial [OS=Homo sapiens] | -0.7 | 0.1 |
| Q92522 | Histone H1x [OS=Homo sapiens] | -0.7 | 0.1 |
| P08754 | Guanine nucleotide-binding protein G(k) subunit alpha [OS=Homo sapiens] | -0.7 | 0.5 |
| Q96EY1-1 | DnaJ homolog subfamily A member 3, mitochondrial [OS=Homo sapiens] | -0.7 | 0.2 |
| P61247 | 40S ribosomal protein S3a [OS=Homo sapiens] | -0.7 | 0.6 |
| P35237 | serpin B6 [OS=Homo sapiens] | -0.8 | 0.2 |
| P12236 | ADP/ATP translocase 3 [OS=Homo sapiens] | -0.8 | 0.6 |
| Q562R1 | Beta-actin-like protein 2 [OS=Homo sapiens] | -0.8 | 0.4 |
| P17858-1 | ATP-dependent 6-phosphofructokinase, liver type [OS=Homo sapiens] | -0.8 | 0.7 |
| Q8IVT2 | Mitotic interactor and substrate of PLK1 [OS=Homo sapiens] | -0.8 | 0.3 |
| Q9HAV0 | Guanine nucleotide-binding protein subunit beta-4 [OS=Homo sapiens] | -0.8 | 0.1 |
| O00763 | Acetyl-CoA carboxylase 2 [OS=Homo sapiens] | -0.9 | 1.1 |
| P49790 | Nuclear pore complex protein Nup153 [OS=Homo sapiens] | -0.9 | 0.2 |
| P51571 | translocon-associated protein subunit delta [OS=Homo sapiens] | -0.9 | 0.2 |
| P04899-4 | Isoform sGi2 of Guanine nucleotide-binding protein G(i) subunit alpha-2 [OS=Homo sapiens] | -0.9 | 1.1 |
| P46777 | 60S ribosomal protein L5 [OS=Homo sapiens] | -1.0 | 0.1 |
| P62879 | Guanine nucleotide-binding protein G(I)/G(S)/G(T) subunit beta-2 [OS=Homo sapiens] | -1.0 | 0.1 |
| Q9Y512 | sorting and assembly machinery component 50 homolog [OS=Homo sapiens] | -1.1 | 0.2 |
| Q6WCQ1-2 | Isoform 2 of Myosin phosphatase Rho-interacting protein [OS=Homo sapiens] | -1.1 | 0.2 |
| Q9Y6I3-1 | Isoform 2 of Epsin-1 [OS=Homo sapiens] | -1.1 | 1.6 |
| P23284 | peptidyl-prolyl cis-trans isomerase B [OS=Homo sapiens] | -1.1 | 0.5 |
| Q5T750 | Skin-specific protein 32 [OS=Homo sapiens] | -1.1 | 0.2 |
| P62263 | 40S ribosomal protein S14 [OS=Homo sapiens] | -1.2 | 0.8 |
| P05166-2 | Isoform 2 of Propionyl-CoA carboxylase beta chain, mitochondrial [OS=Homo sapiens] | -1.2 | 0.9 |
| Q9NZT1 | Calmodulin-like protein 5 [OS=Homo sapiens] | -1.2 | 0.6 |
| P05165-1 | Propionyl-CoA carboxylase alpha chain, mitochondrial [OS=Homo sapiens] | -1.2 | 1.9 |
| P23528 | Cofilin-1 [OS=Homo sapiens] | -1.2 | 0.1 |
| P02545 | Prelamin-A/C [OS=Homo sapiens] | -1.2 | 0.2 |
| P40429 | 60S ribosomal protein L13a [OS=Homo sapiens] | -1.2 | 0.2 |
| P62873 | Guanine nucleotide-binding protein G(I)/G(S)/G(T) subunit beta-1 [OS=Homo sapiens] | -1.4 | 0.2 |
| P62269 | 40S ribosomal protein S18 [OS=Homo sapiens] | -1.4 | 3.4 |
| Q13085-1 | Acetyl-CoA carboxylase 1 [OS=Homo sapiens] | -1.4 | 1.8 |
| P31327-3 | Isoform 3 of Carbamoyl-phosphate synthase [ammonia], mitochondrial [OS=Homo sapiens] | -1.4 | 1.9 |
| P27708 | CAD protein [OS=Homo sapiens] | -1.4 | 1.9 |
| P62917 | 60S ribosomal protein L8 [OS=Homo sapiens] | -1.4 | 0.6 |
| P62987 | Ubiquitin-60S ribosomal protein L40 [OS=Homo sapiens] | -1.4 | 0.7 |
| P62979 | Ubiquitin-40S ribosomal protein S27a [OS=Homo sapiens] | -1.4 | 0.7 |
| P30050-1 | 60S ribosomal protein L12 [OS=Homo sapiens] | -1.4 | 1.6 |
| O94905-1 | Erlin-2 [OS=Homo sapiens] | -1.5 | 0.4 |
| Q00325-1 | Phosphate carrier protein, mitochondrial [OS=Homo sapiens] | -1.5 | 0.8 |
| P63000-2 | Isoform B of Ras-related C3 botulinum toxin substrate 1 [OS=Homo sapiens] | -1.5 | 0.3 |
| P06748 | Nucleophosmin [OS=Homo sapiens] | -1.5 | 0.2 |
| P62280 | 40S ribosomal protein S11 [OS=Homo sapiens] | -1.6 | 1.4 |
| P12004 | proliferating cell nuclear antigen [OS=Homo sapiens] | -1.6 | 1.3 |
| **Q96HS1-1** | **Serine/threonine-protein phosphatase Pgam5, mitochondrial [OS=Homo sapiens]** | **-1.7** | **0.9** |
| **Q06830** | **peroxiredoxin-1 [OS=Homo sapiens]** | **-1.7** | **0.7** |
| **Q8N3V7-1** | **synaptopodin [OS=Homo sapiens]** | **-1.9** | **0.4** |
| **B0I1T2-1** | **unconventional myosin-Ig [OS=Homo sapiens]** | **-1.9** | **0.5** |
| **Q8NFJ5** | **Retinoic acid-induced protein 3 [OS=Homo sapiens]** | **-1.9** | **1.1** |
| **P81605-2** | **Isoform 2 of Dermcidin [OS=Homo sapiens]** | **-1.9** | **0.5** |
| **P35658-3** | **Isoform 3 of Nuclear pore complex protein Nup214 [OS=Homo sapiens]** | **-1.9** | **1.0** |
| **P61626** | **lysozyme c [OS=Homo sapiens]** | **-1.9** | **0.8** |
| **P47756-1** | **F-actin-capping protein subunit beta [OS=Homo sapiens]** | **-1.9** | **0.4** |
| **P07355** | **Annexin A2 [OS=Homo sapiens]** | **-2.0** | **0.6** |
| **P61313-1** | **60S ribosomal protein L15 [OS=Homo sapiens]** | **-2.0** | **0.3** |
| **Q86V48-1** | **Leucine zipper protein 1 [OS=Homo sapiens]** | **-2.0** | **1.2** |
| **P29508** | **Serpin B3 [OS=Homo sapiens]** | **-2.1** | **0.4** |
| **O43707** | **Alpha-actinin-4 [OS=Homo sapiens]** | **-2.1** | **1.5** |
| **Q14978-2** | **Isoform Beta of Nucleolar and coiled-body phosphoprotein 1 [OS=Homo sapiens]** | **-2.1** | **1.2** |
| **Q15149-1** | **plectin [OS=Homo sapiens]** | **-2.2** | **0.8** |
| **Q12904-2** | **Isoform 2 of Aminoacyl tRNA synthase complex-interacting multifunctional protein 1 [OS=Homo sapiens]** | **-2.3** | **0.8** |
| **P51572-2** | **Isoform 2 of B-cell receptor-associated protein 31 [OS=Homo sapiens]** | **-2.3** | **1.4** |
| **Q13263** | **Transcription intermediary factor 1-beta [OS=Homo sapiens]** | **-2.3** | **1.2** |
| **O95425** | **Supervillin [OS=Homo sapiens]** | **-2.3** | **0.6** |
| **P09493-5** | **Isoform 5 of Tropomyosin alpha-1 chain [OS=Homo sapiens]** | **-2.4** | **0.4** |
| **P31153** | **S-adenosylmethionine synthase isoform type-2 [OS=Homo sapiens]** | **-2.4** | **0.6** |
| **O75477** | **erlin-1 [OS=Homo sapiens]** | **-2.4** | **0.7** |
| Q86YZ3 | Hornerin [OS=Homo sapiens] | -2.4 | 1.2 |
| P25311 | Zinc-alpha-2-glycoprotein [OS=Homo sapiens] | -2.5 | 1.1 |
| Q9UHB6-4 | Isoform 4 of LIM domain and actin-binding protein 1 [OS=Homo sapiens] | -2.5 | 0.8 |
| Q9UHB6 | LIM domain and actin-binding protein 1 [OS=Homo sapiens] | -2.5 | 0.8 |
| P04040 | catalase [OS=Homo sapiens] | -2.5 | 0.8 |
| P05109 | Protein S100-A8 [OS=Homo sapiens] | -2.6 | 0.4 |
| P50402 | Emerin [OS=Homo sapiens] | -2.6 | 0.7 |
| P60660 | Myosin light polypeptide 6 [OS=Homo sapiens] | -2.6 | 0.5 |
| Q96P63-2 | Isoform 2 of Serpin B12 [OS=Homo sapiens] | -2.7 | 1.3 |
| Q08554-2 | Isoform 1B of Desmocollin-1 [OS=Homo sapiens] | -2.7 | 0.6 |
| P68032 | Actin, alpha cardiac muscle 1 [OS=Homo sapiens] | -2.7 | 1.0 |
| Q08188 | Protein-glutamine gamma-glutamyltransferase E [OS=Homo sapiens] | -2.8 | 0.7 |
| P60953 | Cell division control protein 42 homolog [OS=Homo sapiens] | -2.8 | 2.7 |
| Q13501-1 | sequestosome-1 [OS=Homo sapiens] | -2.9 | 2.6 |
| P48594 | Serpin B4 [OS=Homo sapiens] | -2.9 | 0.5 |
| O75955 | Flotillin-1 [OS=Homo sapiens] | -2.9 | 0.9 |
| Q96C19 | EF-hand domain-containing protein D2 [OS=Homo sapiens] | -3.0 | 1.1 |
| O14974 | Protein phosphatase 1 regulatory subunit 12A [OS=Homo sapiens] | -3.0 | 1.2 |
| Q02978 | Mitochondrial 2-oxoglutarate/malate carrier protein [OS=Homo sapiens] | -3.0 | 0.8 |
| Q02218 | 2-oxoglutarate dehydrogenase, mitochondrial [OS=Homo sapiens] | -3.0 | 1.0 |
| O14639-5 | Isoform 5 of Actin-binding LIM protein 1 [OS=Homo sapiens] | -3.0 | 0.9 |
| P60174 | Triosephosphate isomerase [OS=Homo sapiens] | -3.1 | 1.2 |
| Q8WWI1-3 | Isoform 3 of LIM domain only protein 7 [OS=Homo sapiens] | -3.1 | 1.8 |
| Q02413 | Desmoglein-1 [OS=Homo sapiens] | -3.1 | 0.7 |
| O75489 | NADH dehydrogenase [ubiquinone]<br>iron-sulfur protein 3, mitochondrial<br>[OS=Homo sapiens] | -3.1 | 4.0 |
| P52907 | F-actin-capping protein subunit alpha-1<br>[OS=Homo sapiens] | -3.2 | 0.6 |
| P60468 | protein transport protein Sec61 subunit<br>beta [OS=Homo sapiens] | -3.2 | 1.1 |
| P32119 | Peroxiredoxin-2 [OS=Homo sapiens] | -3.3 | 0.6 |
| Q15517 | corneodesmosin [OS=Homo sapiens] | -3.4 | 0.5 |
| P11498 | pyruvate carboxylase, mitochondrial<br>[OS=Homo sapiens] | -3.5 | 3.3 |
| P04792 | Heat shock protein beta-1 [OS=Homo<br>sapiens] | -3.5 | 1.5 |
| Q01650 | large neutral amino acids transporter<br>small subunit 1 [OS=Homo sapiens] | -3.6 | 3.0 |
| P15924-1 | Desmoplakin [OS=Homo sapiens] | -3.6 | 0.6 |
| P31944 | Caspase-14 [OS=Homo sapiens] | -3.6 | 2.3 |
| P22735 | Protein-glutamine gamma-glutamyltransferase K [OS=Homo<br>sapiens] | -3.7 | 1.3 |
| Q14847-2 | Isoform 2 of LIM and SH3 domain<br>protein 1 [OS=Homo sapiens] | -3.7 | 4.1 |
| Q5SRD1 | Putative mitochondrial import inner<br>membrane translocase subunit Tim23B<br>[OS=Homo sapiens] | -3.7 | 0.8 |
| P05787-2 | Isoform 2 of Keratin, type II cytoskeletal<br>8 [OS=Homo sapiens] | -3.8 | 0.8 |
| Q13813 | Spectrin alpha chain, non-erythrocytic 1<br>[OS=Homo sapiens] | -4.0 | 1.8 |
| Q13813-2 | Isoform 2 of Spectrin alpha chain, non-erythrocytic 1 [OS=Homo sapiens] | -4.0 | 1.8 |
| Q9ULV4-3 | Isoform 3 of Coronin-1C [OS=Homo<br>sapiens] | -4.0 | 0.9 |
| P52943 | Cysteine-rich protein 2 [OS=Homo<br>sapiens] | -4.0 | 3.1 |
| Q01082-3 | Isoform 2 of Spectrin beta chain, non-erythrocytic 1 [OS=Homo sapiens] | -4.0 | 2.2 |
| Q6NYC8 | Phostensin [OS=Homo sapiens] | -4.2 | 2.4 |
| P50416 | Carnitine O-palmitoyltransferase 1, liver isoform [OS=Homo sapiens] | -4.3 | 3.9 |
| Q01082-1 | Spectrin beta chain, non-erythrocytic 1 [OS=Homo sapiens] | -4.3 | 2.4 |
| O95816 | BAG family molecular chaperone regulator 2 [OS=Homo sapiens] | -4.4 | 2.9 |
| P16403 | Histone H1.2 [OS=Homo sapiens] | -4.5 | 2.0 |
| P10412 | Histone H1.4 [OS=Homo sapiens] | -4.5 | 2.0 |
| P49207 | 60S ribosomal protein L34 [OS=Homo sapiens] | -4.5 | 0.7 |
| Q9NQT5 | exosome complex component RRP40 [OS=Homo sapiens] | -4.5 | 2.6 |
| P06753-2 | Isoform 2 of Tropomyosin alpha-3 chain [OS=Homo sapiens] | -4.6 | 1.7 |
| P50238 | Cysteine-rich protein 1 [OS=Homo sapiens] | -4.6 | 3.7 |
| Q16643-3 | Isoform 3 of Drebrin [OS=Homo sapiens] | -4.7 | 1.3 |
| Q9UDY2-7 | Isoform 7 of Tight junction protein ZO-2 [OS=Homo sapiens] | -4.7 | 3.3 |
| P14923 | Junction plakoglobin [OS=Homo sapiens] | -4.7 | 0.7 |
| P62081 | 40S ribosomal protein S7 [OS=Homo sapiens] | -4.8 | 1.0 |
| P05089-2 | Isoform 2 of Arginase-1 [OS=Homo sapiens] | -5.0 | 1.2 |
| O14950 | Myosin regulatory light chain 12B [OS=Homo sapiens] | -5.3 | 4.1 |
| Q9NYL9 | tropomodulin-3 [OS=Homo sapiens] | -5.4 | 1.4 |
| P46779-3 | Isoform 3 of 60S ribosomal protein L28 [OS=Homo sapiens] | -5.5 | 3.3 |
| P68431 | Histone H3.1 [OS=Homo sapiens] | -5.5 | 1.5 |
| P16401 | Histone H1.5 [OS=Homo sapiens] | -5.7 | 1.8 |
| Q5D862 | Filaggrin-2 [OS=Homo sapiens] | -5.9 | 1.6 |
| O43324-1 | Eukaryotic translation elongation factor 1 epsilon-1 [OS=Homo sapiens] | -6.2 | 3.7 |
| P08670 | Vimentin [OS=Homo sapiens] | -6.4 | 1.2 |
| Q13835 | Plakophilin-1 [OS=Homo sapiens] | -6.6 | 1.3 |
| P62899 | 60S ribosomal protein L31 [OS=Homo sapiens] | -6.7 | 3.8 |
| **P62854** | **40S ribosomal protein S26 [OS=Homo sapiens]** | **-6.7** | **2.2** |
| **P08708** | **40S ribosomal protein S17 [OS=Homo sapiens]** | **-7.1** | **3.7** |
| **P46782** | **40S ribosomal protein S5 [OS=Homo sapiens]** | **-7.3** | **5.0** |
| **Q07157** | **Tight junction protein ZO-1 [OS=Homo sapiens]** | **-7.8** | **3.3** |
| **Q07157-2** | **Isoform Short of Tight junction protein ZO-1 [OS=Homo sapiens]** | **-7.8** | **3.3** |
| **P62841** | **40S ribosomal protein S15 [OS=Homo sapiens]** | **-8.2** | **5.7** |
| **P62633-1** | **Cellular nucleic acid-binding protein [OS=Homo sapiens]** | **-8.5** | **3.4** |
| **P62633-4** | **Isoform 4 of Cellular nucleic acid-binding protein [OS=Homo sapiens]** | **-8.5** | **3.4** |

**Supporting Table S2.**
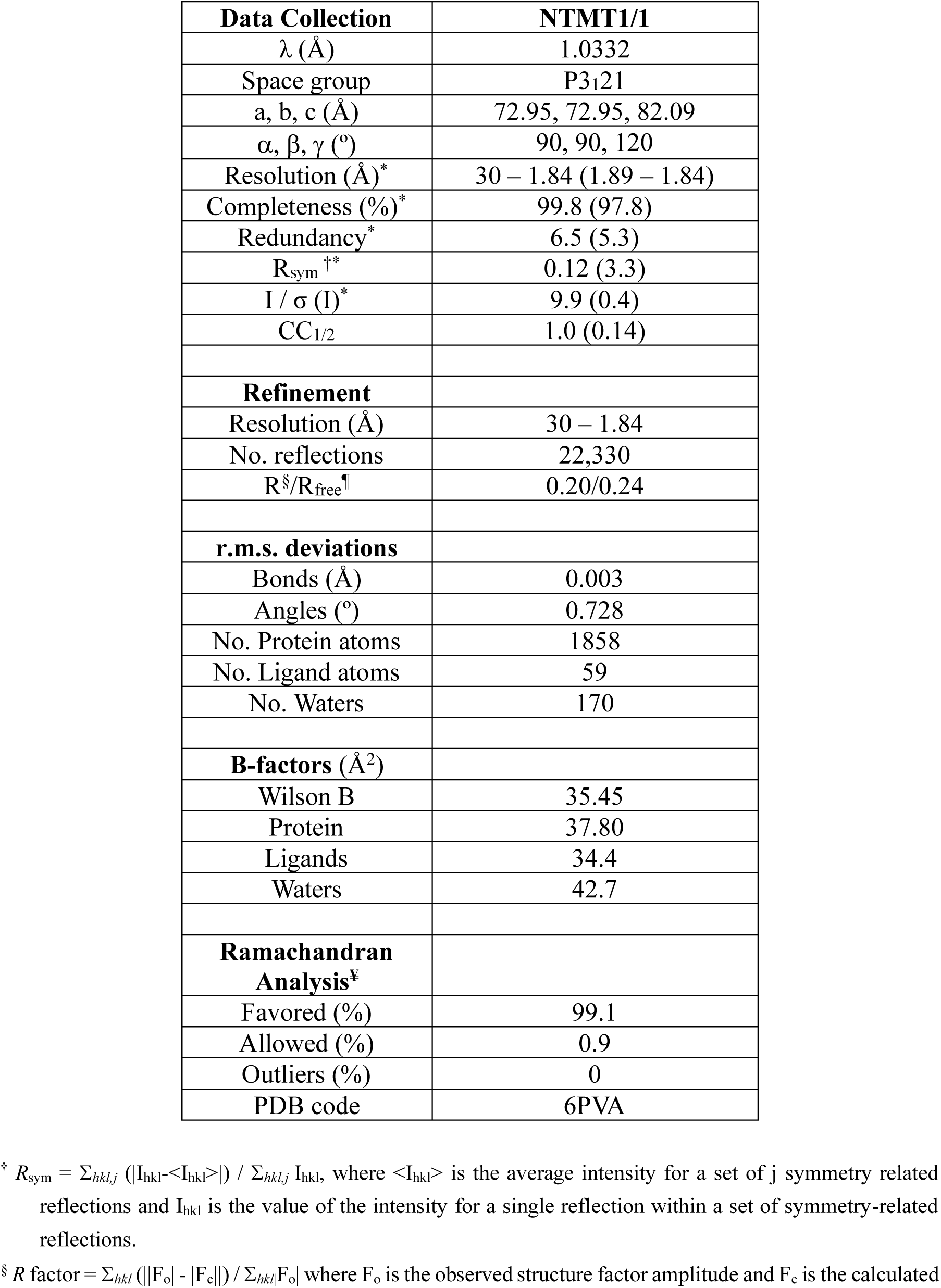
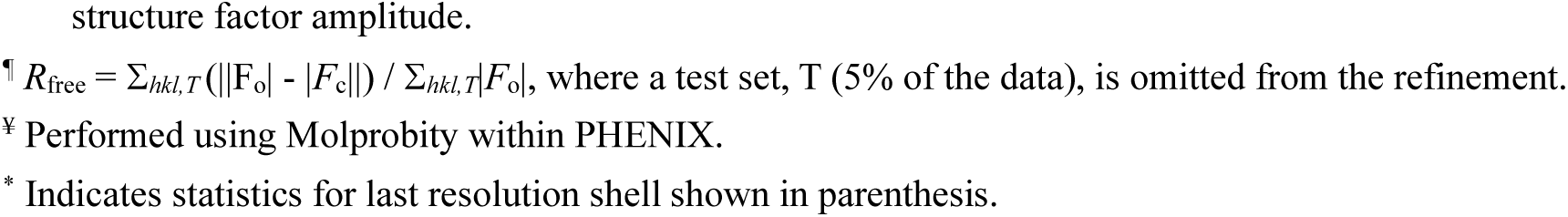
Crystallography data and refinement statistics (PDB ID: 6PVA).

| **Data Collection** | **NTMT1/1** |
| --- | --- |
| $\lambda$ (Å) | 1.0332 |
| Space group | P3<sub>1</sub>21 |
| a, b, c (Å) | 72.95, 72.95, 82.09 |
| $\alpha$, $\beta$, $\gamma$ (°) | 90, 90, 120 |
| Resolution (Å)* | 30 – 1.84 (1.89 – 1.84) |
| Completeness (%)* | 99.8 (97.8) |
| Redundancy* | 6.5 (5.3) |
| $R_{\text{sym}}$†* | 0.12 (3.3) |
| I / $\sigma$ (I)* | 9.9 (0.4) |
| CC<sub>1/2</sub> | 1.0 (0.14) |
| **Refinement** |  |
| Resolution (Å) | 30 – 1.84 |
| No. reflections | 22,330 |
| $R^{\S}/R_{\text{free}}^{\P}$ | 0.20/0.24 |
| **r.m.s. deviations** |  |
| Bonds (Å) | 0.003 |
| Angles (°) | 0.728 |
| No. Protein atoms | 1858 |
| No. Ligand atoms | 59 |
| No. Waters | 170 |
| **B-factors (Å<sup>2</sup>)** |  |
| Wilson B | 35.45 |
| Protein | 37.80 |
| Ligands | 34.4 |
| Waters | 42.7 |
| **Ramachandran Analysis<sup>‡</sup>** |  |
| Favored (%) | 99.1 |
| Allowed (%) | 0.9 |
| Outliers (%) | 0 |
| PDB code | 6PVA |

† $R_{\text{sym}} = \sum_{hkl,j} (|I_{hkl} - \langle I_{hkl} \rangle|) / \sum_{hkl,j} I_{hkl}$ , where $\langle I_{hkl} \rangle$ is the average intensity for a set of j symmetry related reflections and $I_{hkl}$ is the value of the intensity for a single reflection within a set of symmetry-related reflections.
§ $R$ factor = $\sum_{hkl} (||F_o| - |F_c||) / \sum_{hkl} |F_o|$ where $F_o$ is the observed structure factor amplitude and $F_c$ is the calculated
structure factor amplitude.
<sup>¶</sup> $R_{\text{free}} = \sum_{hkl, T} (||F_o| - |F_c||) / \sum_{hkl, T} |F_o|$ , where a test set, T (5% of the data), is omitted from the refinement.
<sup>‡</sup> Performed using Molprobity within PHENIX.
\* Indicates statistics for last resolution shell shown in parenthesis.

## REFERENCES

(1) Luo, M. (2018) Chemical and Biochemical Perspectives of Protein Lysine Methylation. Chem. Rev. 118, 6656–6705.

(2) Kaniskan, H. U., Martini, M. L., Jin, J. (2018) Inhibitors of Protein Methyltransferases and Demethylases. Chem. Rev. 118, 989–1068.

(3) Jarrold, J., Davies, C. C. (2019) PRMTs and Arginine Methylation : Cancer’s Best-Kept Secret? Trends Mol. Med. 25, 993–1009.

(4) Gao, J., Wang, B., Yu, H., Wu, G., Wan, C., Liu, W. (2020) Structural Insight into HEMK2-TRMT112 Mediated Glutamine Methylation. Biochem. J. 477, 3833–3838.

(5) Tooley, C. E. S., Petkowski, J. J., Muratore-Schroeder, T. L., Balsbaugh, J. L., Shabanowitz, J., Sabat, M., Minor, W., Hunt, D. F., Macara, I. G. (2010) NRMT Is an Alpha-N-Methyltransferase That Methylates RCC1 and Retinoblastoma Protein. Nature 466, 1125–1128.

(6) Feng, Y., Hadjikyriacou, A., Clarke, S. G. (2014) Substrate Specificity of Human Protein Arginine Methyltransferase 7 (PRMT7): The Importance of Acidic Residues in the Double e Loop. J. Biol. Chem. 289, 32604–32616.

(7) Del Rizzo, P. A., Trievel, R. C. (2014) Molecular Basis for Substrate Recognition by Lysine Methyltransferases and Demethylases. Biochim. Biophys. Acta-Gene Regul. Mech. 1839, 1404–1415.

(8) Dong, C., Mao, Y., Tempel, W., Qin, S., Li, L., Loppnau, P., Huang, R., Min, J. (2015) Structural Basis for Substrate Recognition by the Human N-Terminal Methyltransferase 1. Genes Dev. 29, 2343–2348.

(9) Jakobsson, M. E., Małecki, J. M., Halabelian, L., Nilges, B. S., Pinto, R., Kudithipudi, S., Munk, S., Davydova, E., Zuhairi, F. R., Arrowsmith, C. H., Jeltsch, A., Leidel, S. A., Olsen, J. V., Falnes, P. (2018) The Dual Methyltransferase METTL13 Targets N Terminus and Lys55 of EEF1A and Modulates Codon-Specific Translation Rates. Nat. Commun. 9, 3411.

(10) Frankel, A., Yadav, N., Lee, J., Branscombe, T. L., Clarke, S., Bedford, M. T. (2002) The Novel Human Protein Arginine N-Methyltransferase PRMT6 Is a Nuclear Enzyme Displaying Unique Substrate Specificity. J. Biol. Chem. 277, 3537–3543.

(11) Liu, X., Wang, L., Li, H., Lu, X., Hu, Y., Yang, X., Huang, C., Gu, D. (2014) Coactivator-Associated Arginine Methyltransferase 1 Targeted by MiR-15a Regulates inflammation in Acute Coronary Syndrome. Atherosclerosis 233, 349–356.

(12) Wang, S. M., Dowhan, D. H., Muscat, G. E. O. (2019) Epigenetic Arginine Methylation in Breast Cancer : Emerging Therapeutic Strategies. J. Mol. Endocrinol. 62, R223–R227.

(13) Tsai, W., Niessen, S., Goebel, N., Yates, J. R., Guccione, E., Montminy, M. (2013) PRMT5 Modulates the Metabolic Response to Fasting Signals. Proc. Natl. Acad. Sci. U.S.A. 110, 8870–8875.

(14) Wu, R., Yue, Y., Zheng, X., Li, H. (2015) Molecular Basis for Histone N-Terminal Methylation by NRMT1. Genes Dev. 29, 2337–2342.

(15) Dai, X., Rulten, S. L., You, C., Caldecott, K. W., Wang, Y. (2015) Identification and Functional Characterizations of N-Terminal α-N-Methylation and Phosphorylation of Serine 461 in Human Poly(ADP-Ribose) Polymerase 3. J. Proteome Res. 14, 2575–2582.

(16) Sathyan, K. M., Fachinetti, D., Foltz, D. R. (2017) α-Amino Trimethylation of CENP-A by NRMT Is Required for Full Recruitment of the Centromere. Nat. Commun. 8, 1–15.

(17) Dai, X., Otake, K., You, C., Cai, Q., Wang, Z., Masumoto, H., Wang, Y. (2013) Identification of Novel α-N-Methylation of CENP-B That Regulates Its Binding to the Centromeric DNA. J. Proteome Res. 12, 4167–4175.

(18) Cai, Q., Fu, L., Wang, Z., Gan, N., Dai, X., Wang, Y. (2014) α-N-Methylation of Damaged DNA-Binding Protein 2 (DDB2) and Its Function in Nucleotide Excision Repair. J. Biol. Chem. 289, 16046–16056.

(19) Bailey, A. O., Panchenko, T., Sathyan, K. M., Petkowski, J. J., Pai, P.-J., Bai, D. L., Russell, D. H., Macara, I. G., Shabanowitz, J., Hunt, D. F., Black, B. E., Foltz, D. R. (2013) Posttranslational Modification of CENP-A Influences the Conformation of Centromeric Chromatin. Proc. Natl. Acad. Sci. 110, 11827–11832.

(20) Bonsignore, L. A., Tooley, J. G., Van Hoose, P. M., Wang, E., Cheng, A., Cole, M. P., Schaner Tooley, C. E. (2015) NRMT1 Knockout Mice Exhibit Phenotypes Associated with Impaired DNA Repair and Premature Aging. Mech. Ageing Dev. 146–148, 42–52.

(21) Bonsignore, L. A., Butler, J. S., Klinge, C. M., Tooley, C. E. S., Schaner Tooley, C. E. (2015) Loss of the N-Terminal Methyltransferase NRMT1 Increases Sensitivity to DNA Damage and Promotes Mammary Oncogenesis. Oncotarget 6, 12248–12263.

(22) Zhang, G., Richardson, S. L., Mao, Y., Huang, R. (2015) Design, Synthesis, and Kinetic Analysis of Potent Protein N-Terminal Methyltransferase 1 Inhibitors.*Org*. Biomol. Chem. 13, 4149–4154.

(23) Zhang, G., Huang, R. (2016) Facile Synthesis of SAM-Peptide Conjugates through Alkyl Linkers Targeting Protein N-Terminal Methyltransferase 1. RSC Adv. 6, 6768–6771.

(24) Chen, D., Dong, G., Noinaj, N., Huang, R. (2019) Discovery of Bisubstrate Inhibitors for Protein N-Terminal Methyltransferase 1. J. Med. Chem. 62, 3773–3779.

(25) Chen, D., Dong, C., Dong, G., Srinivasan, K., Min, J., Noinaj, N., Huang, R. (2020) Probing the Plasticity in the Active Site of Protein N-Terminal Methyltransferase 1 Using Bisubstrate Analogues. J. Med. Chem. 63, 8419–8431.

(26) Metzger, E., Wang, S., Urban, S., Willmann, D., Schmidt, A., Offermann, A., Allen, A., Sum, M., Obier, N., Cottard, F., Ulferts, S., Preca, B. T., Hermann, B., Maurer, J., Greschik, H., Hornung, V., Einsle, O., Perner, S., Imhof, A., Jung, M., Schüle, R. (2019) KMT9 Monomethylates Histone H4 Lysine 12 and Controls Proliferation of Prostate Cancer Cells. Nat. Struct. Mol. Biol. 26, 361–371.

(27) Allali-hassani, A., Wasney, G. A., Siarheyeva, A., Hajian, T., Vedadi, M., Cheryl, H. (2012) Fluorescence-Based Methods for Screening Writers and Readers of Histone Methyl Marks. J. Biomol. Screening 17, 71–84.

(28) Richardson, S. L., Mao, Y., Zhang, G., Hanjra, P., Peterson, D. L., Huang, R. (2015) Kinetic Mechanism of Protein N-Terminal Methyltransferase 1. J. Biol. Chem. 290, 11601–11610.

(29) Dundas, C. M., Demonte, D., Park, S. (2013) Streptavidin-Biotin Technology: Improvements and Innovations in Chemical and Biological Applications. Appl. Microbiol. Biotechnol. 97, 9343–9353.

(30) Woodcock, C. B., Yu, D., Zhang, X., Cheng, X. (2019) Human HemK2/KMT9/N6AMT1 Is an Active Protein Methyltransferase, but Does Not Act on DNA in Vitro, in the Presence of Trm112. Cell Discov. 5, 4–6.

(31) Baysan, M., Woolard, K., Bozdag, S., Riddick, G., Kotliarova, S., Cam, M. C., Belova, G. I., Ahn, S., Zhang, W., Song, H., Walling, J., Stevenson, H., Meltzer, P., Fine, H. A. (2014) Micro-Environment Causes Reversible Changes in DNA Methylation and mRNA Expression Profiles in Patient-Derived Glioma Stem Cells. PLoS One 9, e94045.

(32) Morrison, J. F. (1982) The Slow-Binding and Slow, Tight-Binding Inhibition of Enzyme-Catalysed Reactions. Trends Biochem. Sci. 7, 102–105.

(33) Wu, H., Min, J., Lunin, V. V., Antoshenko, T., Dombrovski, L., Zeng, H., Allali-Hassani, A., Campagna-Slater, V., Vedadi, M., Arrowsmith, C. H., Plotnikov, A. N., Schapira, M. (2010) Structural Biology of Human H3K9 Methyltransferases. PLoS One 5, e8570.

(34) Rizzo, P. A. Del Dirk, L. M. A., Strunk, B. S., Roiko, M. S., Brunzelle, J. S., Houtz, R. L., Trievel, R. C. (2010) SET7 / 9 Catalytic Mutants Reveal the Role of Active Site Water Molecules in Lysine Multiple Methylation. J. Biol. Chem. 285, 31849–31858.

(35) Debler, E. W., Jain, K., Warmack, R. A., Feng, Y., Clarke, S. G., Blobel, G. (2016) A Glutamate / Aspartate Switch Controls Product Specificity in a Protein Arginine Methyltransferase. Proc. Natl. Acad. Sci. U.S.A. 113, 1–6.

(36) Chen, D., Li, L., Diaz, K., Iyamu, I. D., Yadav, R., Noinaj, N., Huang, R. (2019) Novel Propargyl-Linked Bisubstrate Analogues as Tight-Binding Inhibitors for Nicotinamide N-Methyltransferase. J. Med. Chem. 62, 10783–10797.

